# Characterisation of the genetic determinants of context specific DNA methylation in primary monocytes

**DOI:** 10.1101/2023.05.17.541041

**Authors:** James J. Gilchrist, Hai Fang, Sara Danielli, Marketa Tomkova, Isar Nassiri, Esther Ng, Orion Tong, Chelsea Taylor, Hussein Al Mossawi, Evelyn Lau, Matt Neville, Benjamin Schuster-Boeckler, Julian C. Knight, Benjamin P. Fairfax

**Affiliations:** Department of Paediatrics, University of Oxford, Oxford, UK; MRC Weatherall Institute of Molecular Medicine, University of Oxford, Oxford, UK; Wellcome Centre for Human Genetics, University of Oxford, Oxford, UK; Ludwig Cancer Research Oxford, University of Oxford, Oxford, UK; Nuffield Department of Orthopaedics, Rheumatology and Musculoskeletal Sciences, University of Oxford, Oxford, UK; Oxford Centre for Diabetes, Endocrinology and Metabolism, Rheumatology and Musculoskeletal Sciences, University of Oxford, Oxford, UK; Department of Oncology, University of Oxford, Oxford, UK

## Abstract

DNA methylation (DNAm) has pervasive effects on gene expression and associations with ageing-related traits. Here we describe monocyte DNAm responses to inflammatory stimuli across 192 individuals. We find that, unlike the similarly widespread changes in gene expression elicited by LPS and IFN***γ***, DNAm is markedly more sensitive to LPS. Exposure to LPS caused differential methylation at 20,858 immune-modulated CpGs (imCpGs) which display distinct genomic localisation and transcription factor usage, dependent upon whether methylation is lost or gained. Demethylated imCpGs are profoundly enriched for enhancers, and are over-represented by genes implicated in human diseases, most notably cancer. We find LPS-induced demethylation follows hydroxymethylation and for most sites the degree of demethylation correlates with baseline signal. Notably, we find LPS exposure triggers gain in epigenetic age by approximately 6 months, identifying a potential cause of accelerated epigentic aging which has diverse negative health associations. Finally, we explore the effect of genetic variation on LPS-induced changes in DNAm, identifying 209 imCpGs under genetic control. Exploring shared causal loci between LPS-induced DNAm responses and human disease traits highlights examples of human disease associated loci that also modulate imCpG formation.

In summary, our findings suggest innate immune activity continually remodels DNAm in a highly punctate, enhancer-enriched fashion that is under tight genetic control and predominantly involves genes commonly mutated in cancer.

## Introduction

Persistent innate immune activity leads to chronic inflammation, a characteristic of multiple non-communicable disease states, including metabolic syndrome, cardiovascular disease and cancer[1]. DNA methylation (DNAm), the formation of 5-methylcytosine (5mC) at CpG dinucleotides, is a heritable epigenetic modification with profound effects on transcription factor usage and gene expression[2]. Similar to gene expression, DNA methylation is influenced by both local[3] and trans-acting[4] genetic variation forming methyl quantitative trait loci (meQTL). Blood cell DNA methylation is additionally modulated by cellular differentiation, including memory B cell formation[5] and T-cell diversification[6], as well as environmental exposures, most notably smoking[7]. Changes in DNAm accrue with age, an observation that has led to the development of epigenetic clocks, which accurately estimate chronological age[8]. Moreover, age acceleration, as defined by epigenetic aging in excess of chronological age, are associated with health outcomes including longevity[9], cognitive function[10], cancer[11] and all-cause mortality[12]. *Leishmania*[13] and mycobacterial[14] infection, and LPS stimulation in myeloid cells have been shown to induce changes in DNAm. However, how these changes vary between individuals and whether this is influenced by genetic variation is unexplored.

Here, we have addressed how divergent innate immune stimuli modulates DNA methylation in primary monocytes. In a pilot study we found immune stimuli using the Toll-like receptor (TLR) ligands Pam3CysK4, LPS and IFN*γ* have shared and distinct effects on monocyte DNAm. Of the innate stimuli tested, LPS induced the most marked changes in DNAm. We thus explored the effects of LPS stimulation on DNAm at the population level in paired untreated and LPS-treated monocytes from 192 healthy European adults. Using this large cohort we comprehensively define sites of LPS-induced differential methylation. We describe the kinetics of LPS induced DNAm changes, and identify that LPS-induced CpG demethylation is an active process, mediated by ten-eleven translocation (TET) methylcytosine dioxygenases. We demonstrate that LPS stimulation induces changes in the epigenetic clock, with age acceleration equivalent to 6 months seen with 24 hours LPS stimulation. We describe how LPS-induced changes in DNAm vary with respect to genomic location and function, and how baseline DNAm predicts LPS response. Finally, we map genetic determinants of LPS-induced DNAm changes, identifying genetic variation that underlies inter-individual differences in LPS-induced DNAm responses. These genetic predictors of LPS-induced DNAm response colocalise with a range of disease traits, identifying a role for inflammation-induced DNAm changes in human health.

## Results

### TLR agonists and IFN*γ* trigger divergent changes in DNA methylation

To characterise how different immune stimuli affect DNAm, primary monocytes were exposed to either the TLR ligands Pam3CysK4 or LPS, or the Type II interferon IFN*γ*. Pairwise comparison of methylation was analysed in limma[15] alongside incubator controls (24h exposure; n=27 LPS, n=11 Pam3CysK4, n=22 IFN*γ*). Whilst most CpGs remained stable, 2,326/468,025 (0.5%) CpGs demonstrated immune sensitivity in methylation across all stimuli (Table S1). Principal component analysis of these immune-modulated CpGs (hereafter referred to as imCpGs) demonstrated treatments elicited shared and divergent effects (Fig. 1a,b). Restricting analysis to samples with stimuli across all conditions (n=11), demonstrated LPS elicited the most detectable changes in methylation, with substantial overlap with PAM3CysK4 responses (Fig. 1c).

**Fig. 1.**
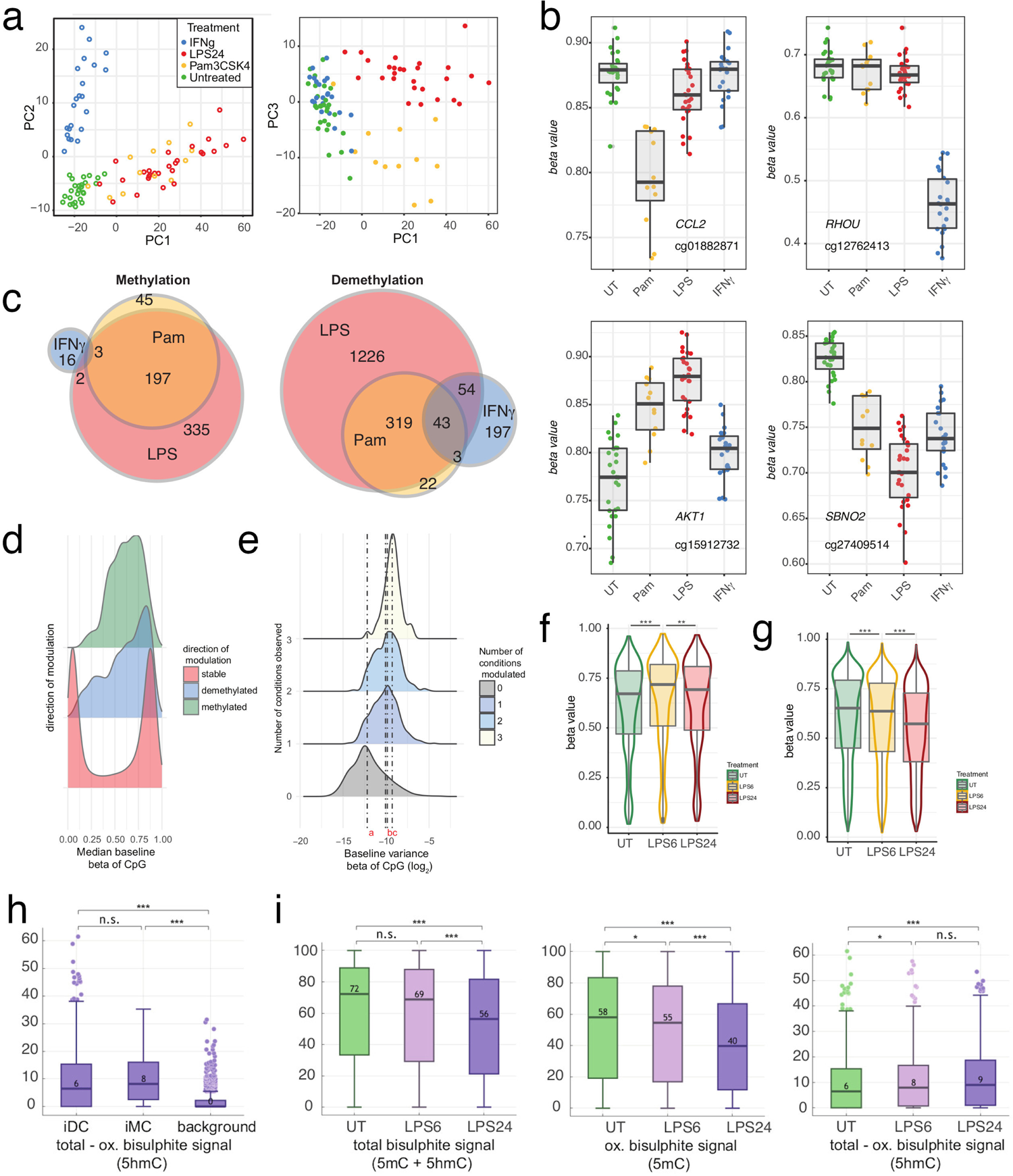
**a** Principal component analysis of 1,134 differentially methylated CpGs across any treatment(FDR *<* 0.05) shows samples cluster according to treatment. **b** Examples of CpGs showing significant differential methylation with divergent treatments including cg01882871 6.2kb upstream of CCL2 (PAM3CysK4 specific demethylation), cg12762413 within the gene body of RHOU (IFN*γ* specific demethylation), cg15912732 within the gene body of AKT1 (PAM3CysK4 and LPS methylation) and cg27409514 intronic to SBNO2 (pan-treatment demethylation). **c** Differentially methylated CpGs seen for each treatment (n=11 individuals) summarised in Venn diagrams. **d** Density plot of median untreated beta of CpGs from pilot samples comparing CpGs that are stable versus those that are either methylated or demethylated across any condition. **e** Variance of beta values for a CpG in untreated samples according to the number of treatments under which it is observed to be modulated. CpGs not observed to change with treatment have reduced variance across the cohort compared to those modulated in one condition (*p <* 2.2 *×* 10*^−^*^16^ wilcoxon rank sum test), 1 vs. 2 conditions (*p* = 0.003) and 2 vs. 3 conditions (*p* = 0.006). Boxplots of all CpGs **f** significantly methylated with either 6 or 24h LPS or **g** significantly demethylated at either 6 or 24h of LPS. \*\**p* = 4.4 *×* 10*^−^*^4^,\*\*\**p <* 2.2 *×* 10*^−^*^16^ (Wilcoxon tests of either UT vs. 6h LPS or 6h LPS versus 24h LPS). **h** Baseline percentage 5hmC methylation at demethylated immune-modulated CpGs (iDC), methylated immune-modulated CpGs (iMC) and across all sequenced CpGs (background), as assayed with paired bisulfite (BS) and oxidative bisulphite (OxBS) sequencing. **i** Percentage total (left), 5mC (middle) and 5hmC (right) methylation at LPS-demethylated immune-modulated CpGs in the untreated state and following 6 and 24 hours of LPS stimulation, as assayed with paired BS and OxBS sequencing.

Similar to CpGs that vary across cell type [16], imCpGs had comparatively more intermediate methylation compared to stable CpGs in the untreated state (*p <* 2.2 *×* 10*^−^*^16^, Kolgomorov-Smirnov test, Fig. 1d) and demonstrated elevated baseline variance in methylation between individuals. This increased with the number of conditions the CpG was sensitive to (Fig. 1e) and suggests that a subset of monocyte CpGs are primed to innate-immune stimuli. To explore the effect of time on LPS-modulated DNAm, we treated monocytes with LPS for 6 or 24h (n=18). We found LPS induced gain in methylation was an early phenomenon; with 126 and 135 imCpGs showing increased methylation signal at 6h and 24h respectively (Table S2). On average, CpGs with increased methylation signal at 24h showed more marked methylation induction at 6h, indicating maximal gain in methylation occurs early with subsequent demethylation at these sites (Fig. 1f, Fig. S1). Notably, 11 imCpGs showed increased beta values at 6h with a net loss in beta values detected by 24h (Fig. S1). Conversely, 99.4% (1281/1289) demethylated ImCpGs at 6h demonstrated further demethylation by 24h, indicating demethylation is more continuous and prolonged (Fig. 1g, Fig. S1, Table S2).

### LPS induced demethylation involves the formation of 5-hmC

CFSE cell proliferation assays confirmed LPS did not induce monocyte cell division under the conditions used, indicating LPS observed demethylation is active. Active demethylation involves TET enzymes that catalyse the oxidation of 5mC to form 5-hydroxymethylcytosine (5-hmC)[17, 18] with successive oxidation culminating in base-excision repair and replacement with non-methylated cytosine. To ascertain if LPS-induced demethylation involved 5-hmC formation in monocytes, we used pairwise bisulfite (BS) and oxidative bisulfite (OxBS) sequencing[19] of target-captured methylome DNA from 4 individual monocyte samples at baseline, 6 and 24h of LPS treatment. Across pooled reads from all samples we identified 107 imCpGs identified in the array timecourse experiment where we had sufficient coverage to monitor methylation in the sequencing data. ImCpGs have significantly higher baseline 5hmC levels compared to background CpGs similarly covered in arrays (iDCs 6%, iMCs 8%, vs 0.5% background *p <* 0.001) (Figure 1h). We note a fall in both total and 5mC signals at array determined LPS demethylated imCpGs at 6 and 24h (Fig. 1i), whereas the 5hmC signal at these sites increases at both 6 and 24h, demonstrating that LPS induced demethylation involves the formation of this intermediary (Fig. 1i).

### Differential methylation in response to LPS across a population

To explore variation in observed changes in methylation across a population of individuals we focused on the 24h LPS response. We increased the total sample size to 192. After quality control, we generated paired untreated vs. LPS methylation data for a total of 188 individuals. LPS stimulation modulated DNAm at 20,858 imCpGs (5.1% of tested, *FDR <* 0.01), with 10,614 (2.6%) showing gain in methylation and 10,244 (2.5%) demethylation (Fig. 2a, Table S3). There was high concordance in responses from the original 27 samples and the further 161 samples (*p <* 2.2 *×* 10*^−^*^16^, Fig. S2). We identified differentially methylated regions (DMRs) induced by LPS with DMRcate[20], observing 6,473 regions where two or more contiguous CpGs showed differential methylation (50.8% demethylated, median length 5 CpGs, range:2-94 CpGs, 49.2% of which were methylated, median length 4 CpGs, range:2-82 CpGs, Kolmogorov Smirnov test *p* = 0.0003, Table S4). Highly demethylated regions were observed across the genes *DAXX*, *CUX1* and *ARID5B*, with the most significant DMR in *DAXX*, whilst the most demethylated CpGs were observed in the eighth intron of the gene *CUX1* (maximally at cg15755348), encoding a protein with transcriptional repressor and activator properties (CUX1, cut-like homeobox 1)[21] frequently haploinsufficent or mutated across haematological malignancies and solid tumours[22, 23]. Notably, this region was the most demethylated using BS and OxBS sequencing (Fig. S3). We also noted demthylation across multiple interleukin encoding genes and receptors including *IL36RG*, *IL1RN*, *IL1A* and *IL7R*, which we have previously shown to be strongly induced in monocytes by LPS[24]. Examples of regions gaining methylation signal include *CHID1*, *AKT1* and *ITGAE*. With the largest induction being noted at cg15912732 in the third intron of *AKT1*. *AKT1* encodes Protein Kinase B (PKB) which is sentinel to cell growth and division. Interestingly, deletion of *CUX1*, which is antagonistic to the PI3K-1 pathway, results in hyper-activation of AKT1[23] suggesting synergistic effects in the changes observed. Given the number of DMRs defined over the cohort varies according to the parameters provided to the model, and the relative sparsity in coverage from the array, we focused downstream analysis on single CpGs.

**Fig. 2.**
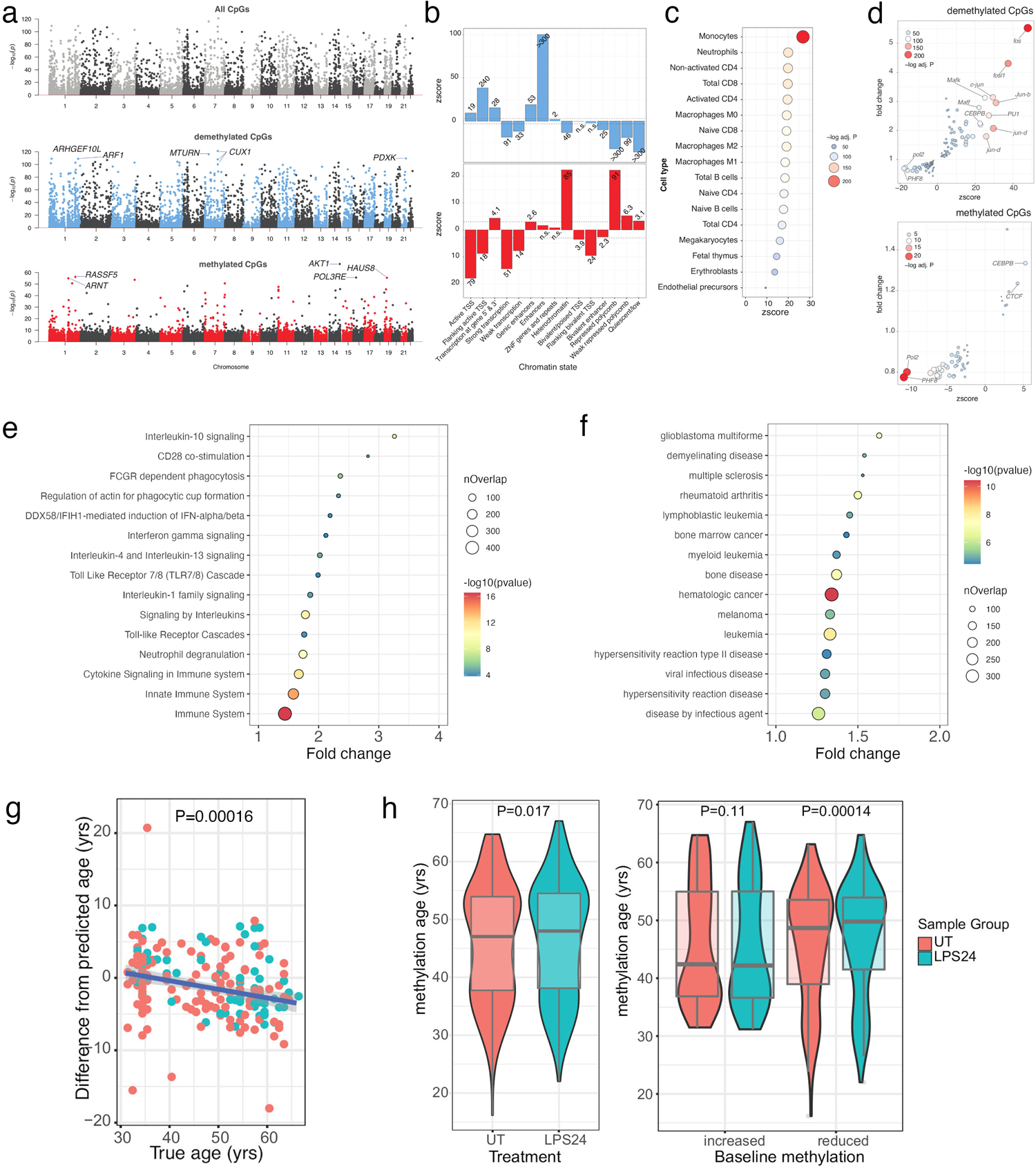
**a** Manhattan plots of differentially methylated CpGs in response to LPS stimulation; all CpGs (top), demethylated CpGs (blue, middle) and CpGs with gains of methylation (red, bottom). **b** Enrichment of significantly demethylated imCpGs (blue, top) and CpGs with significant gains of methylation in response to LPS stimulation (red, bottom) with ENCODE chromatin state segmentations. **c** Enrichment of demethylated imCpGs within promoter-captured enhancer DNA regions across 17 cell types. **d** Enrichment of imCpGs within transcription factor binding sites in K562 cells. Reactome Immune System (**e**) and Disease Ontology (**f**) pathway analysis of genes proximal (with 5kb) to LPS-demethylated CpGs in monocytes. All depicted pathways are significantly enriched (adjusted *p <* 0.05), with the most signficantly enriched pathways from each ontology plotted; Reactome Immune System (15 of 57), Disease Ontology (15 of 86). nOverlap, number of overlapping genes from a pathway. Enrichment is calculated using Fisher Exact tests. **g** Correlation of methylation age vs chronological age for monocytes (untreated) from 92 individuals (male, red; female, blue). **h** Effect of LPS stimulation on methylation age (left). Relationship between baseline methylation age vs chronological age (individuals dichotomised as methylation increased or reduced with respect to chronological age) and change in methylation age on LPS stimulation (right).

### Genomic organisation of differential methylation

We assessed the distribution of imCpGs among chromatin state annotations in primary monocytes[25] (Fig. 2b). We found imCpGs show enrichment across divergent genomic locations according to their behaviour, with profound enrichment for demethylated imCpGs in enhancer (*p <* 1 *×* 10*^−^*^300^), transcription start site (TSS) flanking (*p* = 1 *×* 10*^−^*^240^), genic enhancer (*p* = 1 *×* 10*^−^*^53^), transcription flanking (*p* = 1 *×* 10*^−^*^28^) and active TSS (*p* = 1 *×* 10*^−^*^19^) regions. The majority of other regions were depleted for demethylated imCpGs, most notably repressed polycomb and quiescent regions (*p <* 1 *×* 10*^−^*^300^). In contrast, imCpGs showing gain in methylation were enriched in heterochromatin and repressed polycomb regions (*p* = 1 *×* 10*^−^*^84^, *p* = 1 *×* 10*^−^*^90^), whereas active TSS, TSS flanking regions and both strongly and weakly transcriptionally active regions were significantly depleted (*p* = 1 *×* 10*^−^*^79^, *p* = 1 *×* 10*^−^*^17^, *p* = 1*×*10*^−^*^50^ & *p* = 1*×*10*^−^*^13^). To ascertain whether imCpGs involved genomic regions with evidence of physical interaction with promoters consistent with enhancer activity we interrogated promoter capture HiC data for 17 different cell types [26]. This demonstrated demethylated imCpGs are highly enriched within promoter-captured enhancer DNA from monocytes (2.4 fold-change, *FDR* = 4.2 *×* 10*^−^*^109^, Fig. 2c). We found no enrichment with methylated imCpGs. In keeping with enhancer usage being influenced by cell-type, the degree of enrichment of demethylated imCpGs to promoter captured DNA across monocytes was markedly greater than the other 16 cell types (Fig. 2c).

### Transcription factor usage

Among transcription factor binding sites (TFBS) in K562 cells, demethylated imCpGs were highly enriched in TFBS of 49 different transcription factors (Fig. 2d, Table S5) with AP-1 subunits showing the most prominent enrichment (FOSL1 5.5x increase, *p* = 4.8 *×* 10*^−^*^247^, JUND 2.1x increase, *p* = 3.2 *×* 10*^−^*^154^). Conversely, there was comparatively limited enrichment for methylated imCpGs overlapping TFBS (Table S6), with subtle enrichment noted for CEBPB and the insulating factor CTCF, in keeping with preferential polycomb location of methylated imCpGs. Similarly, in keeping with enrichment of methylated imCpGs in transcriptionally inactive regions, we observed depletion of multiple TFBS for methylated imCpGs (Fig. 2e, Table S6). The K562 ChIP-seq data does not contain information regarding a number of transcription factors, notably NF-KB which is a key mediator of the LPS response. We therefore also looked at data from the lymphoblastoid cell line GM12878 and observed marked enrichment of demethylated imCpGs (Tables S7 & S8, Fig. S4) for JUN (JUND 7.4x enrichment, *p* = 5.2 *×* 10*^−^*^43^), ATF (BATF 4.2x enrichment, *p* = 6.1 *×* 10*^−^*^232^) and NF-KB (2.1x enrichment *p* = 8.7 *×* 10*^−^*^93^).

### LPS modulated CpGs are informative of human disease

To understand the biological significance of imCpGs we performed enrichment analysis of the most proximal gene (within 5kb window) to an imCpG, using Reactome Immune System[27] (R-HSA-168256) and Disease Ontology[28] gene annotations in XGR[29]. Among methylated imCpGs, of which 5,246 are within 5kb of a gene, there was no evidence for biological pathway or human disease enrichment. By contrast, among genes proximal to demethylated imCpGs (n=5,607), we observed striking enrichment (Fig. 2e, Table S9) for immune-related biological pathways (n=67, *FDR <* 0.05) including phagocytosis (n=3 pathways), TLR signalling (n=19), inflammasome activation (n=3) and cytokine signalling (n=19). We further observed marked enrichment of genes proximal to demethylated imCpGs (Fig. 2f, Table S10) among pathways implicated in the pathogenesis of cancer (n=45 pathways), in particular haematological malignancy (n=13), autoimmune disease (n=8) and viral infection (n=5).

### Treatment with LPS causes epigenetic age acceleration

We next explored the degree to which LPS-induced differential DNAm was associated with age, sex and smoking status by testing for interactions with treatment. We did not observe any CpGs where the degree of differential methylation interacted with these variables, suggesting the effect of LPS was independent of key demographic factors. Whilst the association between DNAm and chronological age is well characterized[30–32], with disease states associated with epigenetic age-acceleration[9–12], the extent to which age-associated CpG methylation is plastic in the short term is unclear. Interestingly, CpGs associated with age acceleration are implicated in TLR and interferon responses[33]. To explore whether treatment with LPS influenced the epigenetic age of the cohort we used the epigenetic clock as described by Horvath to determine whether exposure to LPS caused age acceleration in monocytes[34]. We explored age-related changes in methylation across UT and LPS-treated samples, observing 29,134 (Table S11) and 24,389 (Table S12) CpGs associated with age respectively. There was a high concordance between significant results (*r* = 0.99, *p <* 2.2 *×* 10*^−^*^16^, Fig. S5), with the reduced number of observations after LPS treatment likely reflecting inter-individual variation of LPS responses. Similar to others[35], we observed the clock to significantly underestimate the age of older individuals (*p* = 5.2 *×* 10*^−^*^7^, 95% CI 1.5-3.2 years, n=92, Fig. 2g). Irrespective, this observation should not impact a pairwise comparison of untreated and LPS treated monocytes. This demonstrated that LPS elicited a small but significant increase in epigenetic age across the group (+0.5 yr, 95 CI 0.09-0.91yr, P=0.014, Fig. 2h). Further analysis of this LPS-induced age acceleration showed the effect was confined to those who showed no evidence of baseline epigenetic age acceleration, with the effect of LPS on these individuals leading to a pronounced age acceleration (mean=+1.08, 95% CI:0.54-1.62, *p* = 1.4*×*10*−*4, Fig. 2h). Younger individuals are more likely to have baseline epigenetic age acceleration, and we therefore tested if this result could be confounded by age. This is not the case, with LPS-induced epigenetic age acceleration being restricted to individuals with reduced baseline epigenetic age in both younger and older individuals (Fig. S6). To our knowledge, this observation is the first demonstration of short-term plasticity in the DNAm epigenetic clock, with acceleration being induced by activation of the TLR4 pathway. The restriction of this effect to those without baseline epigenetic age acceleration rules out non-specific sequelae of LPS, and suggests individuals with epigenetic age equal or greater than chronological age have already accrued modifications at these sites through life-events.

### Baseline determinants of change in methylation

Just as environmental stimuli may lead to long-lasting effects on DNAm, we hypothesized that the level of baseline methylation might dictate the propensity to environment-associated modulation. We thus explored whether the degree of baseline methylation of an imCpG altered its sensitivity to LPS changes. For the majority of LPS-modulated imCpGs we find magnitude of response to be highly correlated with baseline methylation (Fig. 3a,b). In most cases, baseline correlated (BC) methylated imCpGs with reduced baseline methylation had larger LPS-induced methylation changes. For demethylated imCpGs, high baseline methylation tended to enhance the degree of demethylation. For 764/10,244 (6.0%) of demethylated imCpGs and 881/10,614 (8.3%) of methylated imCpGs we observe no correlation (*p >* 0.05) between baseline methylation and response to LPS (Fig. 3a,b). Baseline independent (BI) imCpGs demonstrated a significant difference (*p <* 2.2*e−*16) in the distribution of untreated DNAm beta values, with more intermediate baseline methylation values, indicating heterogeneity at these positions amongst cells (Fig. 3c) and suggesting these represent a different subset of LPS-responsive CpGs. We find BI demethylated imCpGs tend to be more responsive to LPS than BC demethylated imCpGs, whereas the effect of LPS on BI methylated imCpGs is less than at BC methylated imCpGs (Fig. 3c). Among imCpGs, BI and BC CpGs diverge in their transcription factor usage (Table S13), for example both BC demethylated imCpGs and methylated imCpGs are highly enriched for ERG1 binding sites (Fig. 3d). ERG1 has been shown to recruit the Ten-Eleven Translocation (TET) enzyme, TET1[36]. This is complementary to our observation that imCpgs are enriched for 5hmC at baseline (Fig. 1h), suggesting a model in which BC imCpGs are primed for LPS-induced methylation changes, in part through ERG1-mediated TET enzyme recruitment and formation of 5hmC.

**Fig. 3.**
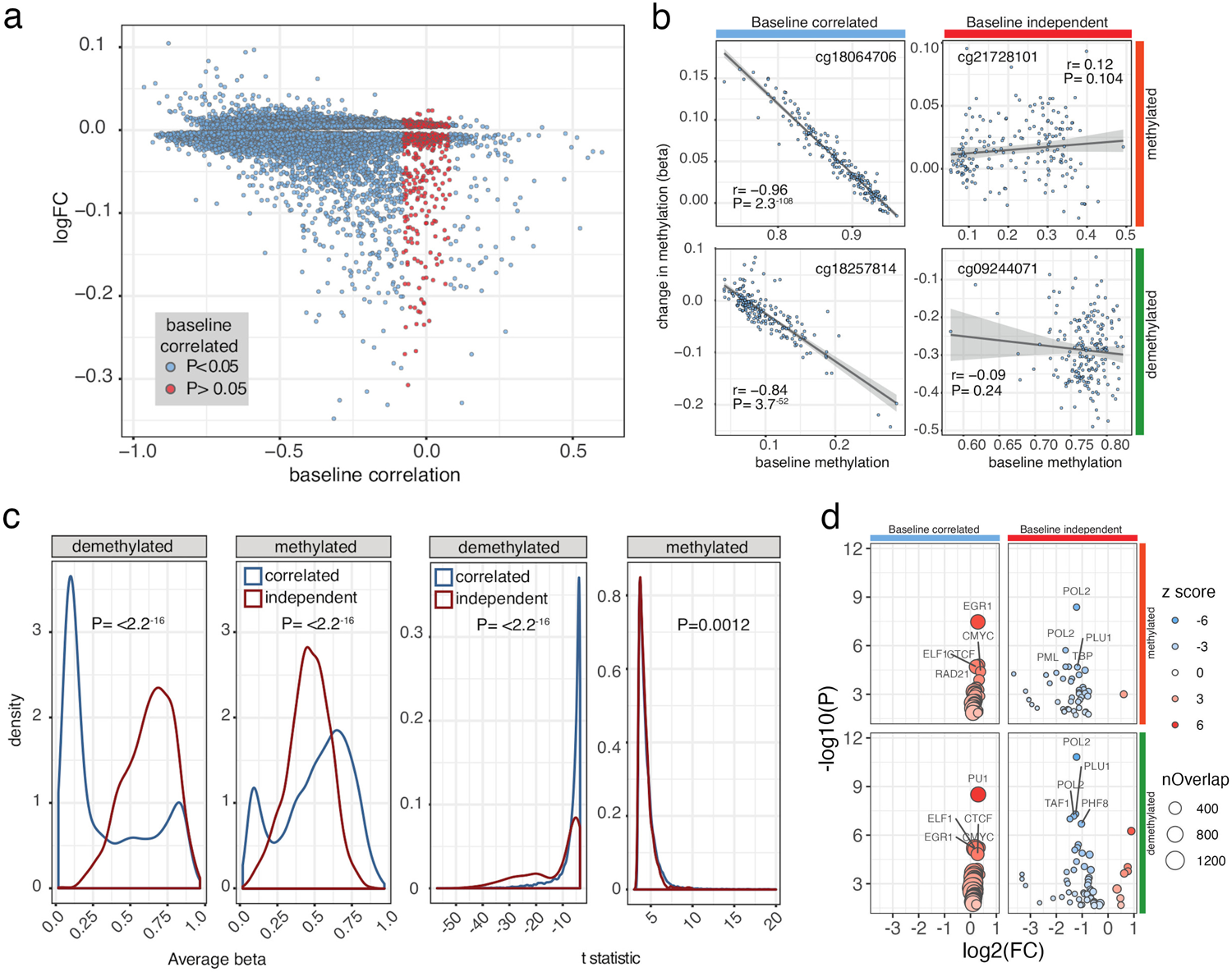
Correlates of differential CpG methylation in response to LPS. **a** Relationship between baseline methylation and change in methylation induced by LPS for all imCpGs. **b** Examples of baseline correlated and baseline independent imCpGs for LPS-methylated and LPS-demethylated loci. Baseline correlated and independent imCpGS have distinct baseline methylation distributions (**c**, left panels) and distinct methylation responses to LPS stimulation (**c**, right panels). **d** Enrichment for overlap with transcription factor binding sites in K562 cells at baseline correlated and independent imCpGs.

### Genetic determinants of LPS-induced methylation changes

Genetic variation is a significant determinant of CpG methylation[3, 4]. In keeping with this, we were able to identify (*FDR <* 0.05) 69,370 and 69,503 mQTL in untreated monocytes and LPS-stimulated monocytes respectively (Table S14 and S15). Given the marked alterations in DNAm observed following LPS stimulation however, we were interested in defining genetic determinants of LPS associated imCpG formation. In a paired analysis (n=186 individuals), we correlated the magnitude of change in methylation on LPS stimulation at the 20,858 imCpGs with genotype at common, well-imputed SNPs within 100kb of the CpG. Mapping immune-modulated methylation quantitative trait loci (im-mQTL) in this way reveals 209 CpG for which changes in methylation in response to LPS stimulation are influenced by germline genetic variation (Fig. 4a, Table S16). We find im-mQTL are more frequent among demethylated imCpGs as compared to methylated imCpGs (186 vs. 23, *p* = 1.46*×*10*^−^*^34^, *OR* = 8.54 Fisher exact test). Complementary to our observation that LPS-demethylated imCpGs are enriched for overlap with AP-1 transcription factor binding sites, im-mQTL at demethylated imCpGs are enriched for Fosl1 binding sites (*p* = 5.0 *×* 10*^−^*^4^, OR 4.13, Fig. 4b, Table S17). We found no evidence of transcription factor binding site enrichment among im-mQTL at methylated imCpGs.

**Fig. 4.**
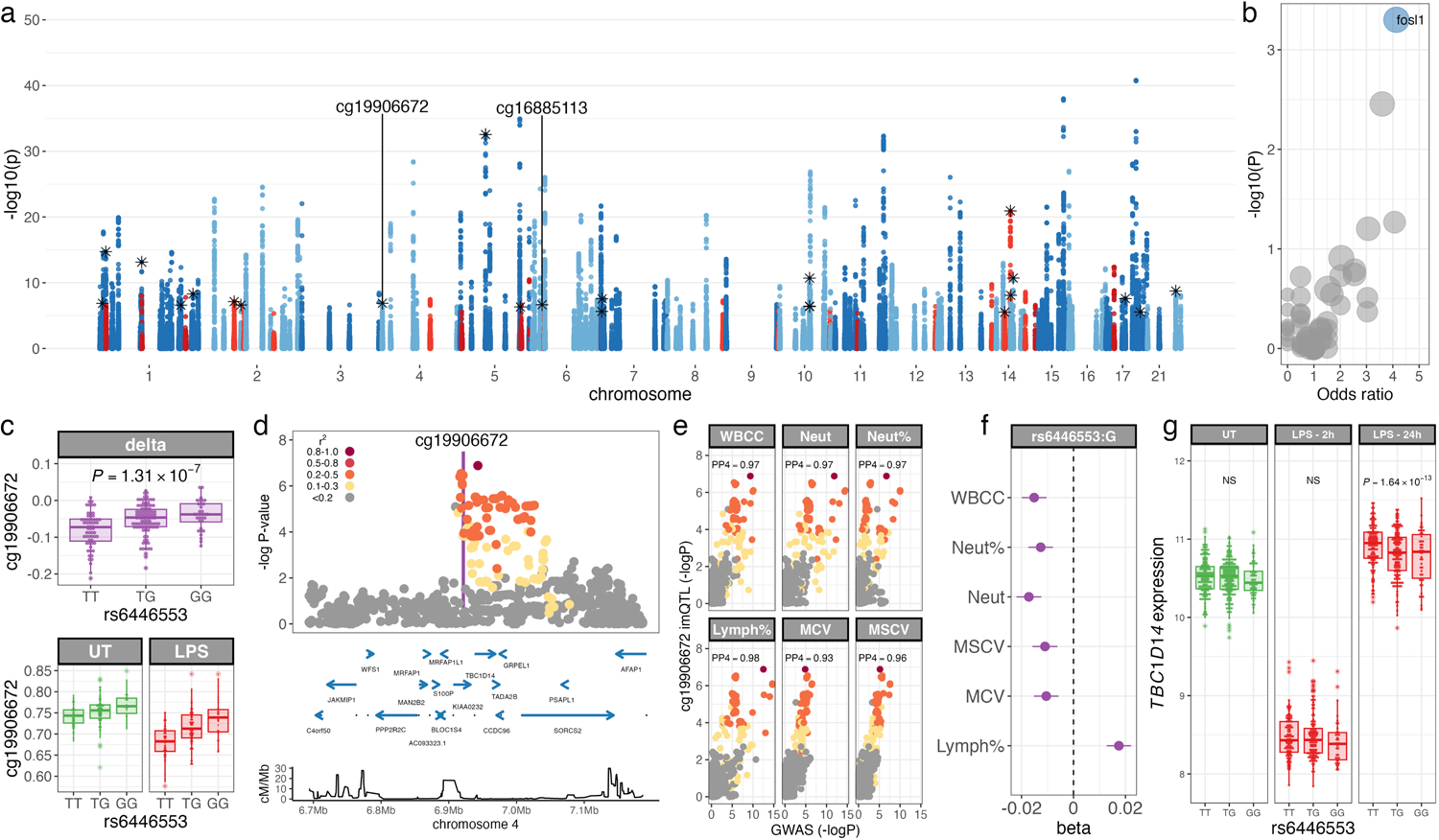
**a** Manhattan plot of immune-modulated methylation quantitative trait loci (im-meQTL) in LPS-stimulated monocytes (n=209). QTL mapping of normalised differential methylation following LPS stimulation in *cis* (within 100kb of the CpG). Im-meQTL are coloured according to effect of LPS on CpG methylation (demethylation, n=186, blue; increased methylation, n=23, red). Im-meQTL (n=23) with evidence of colocalisation to GWAS traits are highlighted (stars). **b** Enrichment of transcription factor binding sites for im-meQTL at demethylated CpGs. Each point represents a distinct transcription factor binding site in ENCODE K562 data (n=83), and point-size is proportional to the number of overlapping im-meQTL with each site. P-values are calculated by permutation. Significantly enriched transcription factors are highlighted (blue). **c** Im-meQTL at cg19906672; correlation of rs6446553 genotype with delta beta methylation (top), and untreated (UT) and LPS-stimulated cg19906672 methylation (bottom). P-values are calculated by linear regression. **d** Regional association plot of cg19906672 im-meQTL. Protein-coding genes are highlighted (blue). Lower panel depicts recombination rate. **e** Colocalisation plots of the cg19906672 im-meQTL with white blood cell count (WBCC), neutrophil count (Neut), neutrophil % (Neut%), lymphocyte % (Lymph%), mean corpuscular volume (MCV) and mean sphered corpuscular volume (MSCV) GWAS. **f** Forest plot depicting effect estimates and 95% confidence intervals of rs6446553:G carriage WBCC, neutrophil count and %, lymphocyte %, MCV and MSCV. **g** Correlation of rs6446553 genotype with *TBC1D14* expression in unstimulated monocytes (green) and following 2 and 24 hours of LPS stimulation (red). NS, not significant. For regional association and colocalisation plots, SNPs are coloured according to strength of LD (CEU population) to the peak im-meQTL SNP. Box and whisker plots; boxes depict the upper and lower quartiles of the data, and whiskers depict the range of the data excluding outliers (outliers are defined as data-points *>* 1.5*×* the inter-quartile range from the upper or lower quartiles).

We explored the relationship between im-mQTL and genetic determinants of human phenotypic variation, observing 23 of 209 im-mQTL to colocalise with a genetic determinant of at least one GWAS trait (Figure 4a, Table S18). An illustrative example of this is cg19906672, which is demethylated in response to LPS, the magnitude of which is influenced by allelic variation at rs6446553, 24Kb downstream from the CpG (*p* = 1.31 *×* 10*^−^*^7^, Fig. 4c,d), with carriers of the T allele showing increased LPS-induced demethylation. This polymorphism colocalises with genetic loci determining several haematological parameters (Fig. 4e,f) including white cell count (posterior probability of colocalisation, PP4=0.97), neutrophil count (PP4=0.97), neutrophil percentage (PP4=0.97), lymphocyte percentage (PP4=0.98), mean corpuscular volume (PP4=0.93) and mean sphered corpuscular volume (PP4=0.96).

Given the variable effects of DNAm on gene expression[37], we sought to identify examples of colocalisation between im-mQTL and the genetic determinants of differential gene expression in LPS-stimulated monocytes in previously-published data[38]. We extracted *cis*-eQTL mapping data for genes within 1Mb of each im-mQTL in unstimulated monocytes and in monocytes following 2 and 24 hours of LPS stimulation. We then used moloc[39] to assess evidence for shared causal loci between each im-mQTL and local eQTL in monocytes across the three stimulation conditions. That analysis supports sharing (*PP >* 0.8) of a causal locus between 13 im-mQTL and at least one *cis* eQTL in monocytes, with evidence of colocalisation at 21 im-mQTL:genic eQTL pairs (Table S19). We identified 4 instances where the genetic determinants of differential methylation on LPS stimulation are shared with regulatory variation controlling gene expression in the untreated state alone, and 12 instances where sharing is only evident following LPS stimulation (Table S19).

The im-mQTL at cg19906672 (rs6446553), as described above, is an example of the latter, with the im-mQTL colocalising with a *cis* eQTL for *TBC1D14* expression only after 24 hours of LPS stimulation. We find that in addition to being associated with increased demethylation, rs6446553:T allele is associated with *TBC1D14* expression post-LPS (Fig. 4g). *TBC1D14* encodes a TBC (Tre-Bub-CDC16) domain-containing GTPase activating protein, which functions as a negative regulator of autophagy[40]. Autophagy is critical to the maintenance of haematopoietic stem cell function, and in keeping with a role for *TBC1D14* expression in myeloid and lymphoid haematological parameters, changes in autophagy are associated with perturbed myeloid:lymphoid ratios[41].

A further example of an im-mQTL informative of human disease is the regulation of cg16885113 LPS-induced methylation by rs3129058 (Fig. 5a,b), which colocalises with a GWAS risk locus for lung cancer[42] (PP4=0.94, Fig. 5c). This im-mQTL also colocalises with an eQTL for expression of *ZFP57*, but only at baseline, i.e. in untreated monocytes (Fig. 5d). We have previously identified an eQTL for *ZFP57* RNA expression in PBMCs at rs375984, and eQTL mapping of regulatory determinants of *ZFP57* RNA expression in naive monocytes (Fig. 5e) demonstrates that rs375984, rs3129058 (the lead im-mQTL variant) and rs417764 (the lead *ZFP57* eSNP in monocytes) are all highly associated with *ZFP57* expression and are in linkage dysequilibrium (*r*^2^ = 0.9, 1000 Genomes Project GBR) with one another. *ZFP57* encodes a Kruppel-associated box containing zinc-finger protein, which is required to maintain DNAm at imprinting control regions[43–45]. We therefore hypothesised that rs417764, as a regulator of *ZFP57* expression, could have genome-wide effects on CpG methylation in monocytes. Of 407,951 CpGs tested, methylation at 151, clustered in 20 genomic regions, is significantly associated (*FDR <* 0.05) with rs417764 genotype (Table S20). In keeping with the established biology of ZFP57, these CpGs are highly enriched (*p* = 4.5 *×* 10*^−^*^49^, OR 75.75) for known imprinted regions[46–50] (Table S21), which cluster in two genomic regions; chr6:29,648,344-29,649,133 and chr20:57,426,997-57,427,972 (Fig. 5f).

**Fig. 5.**
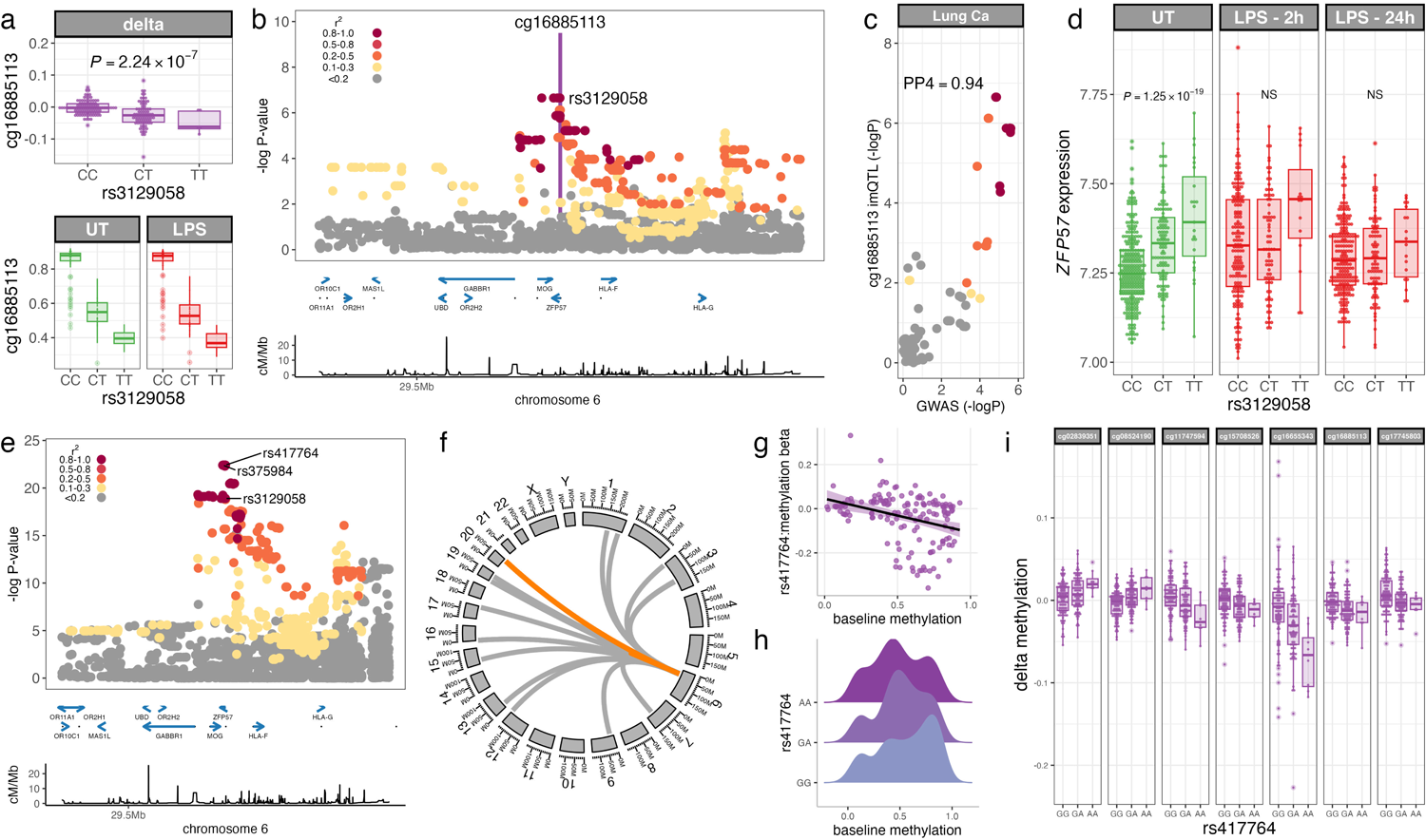
**a** Im-meQTL at cg16885113; correlation of rs3129058 genotype with delta beta methylation (top), and untreated (UT) and LPS-stimulated cg16885113 methylation (bottom). P-values are calculated by linear regression. **b** Regional association plot of cg16885113 im-meQTL. Protein-coding genes are highlighted (blue). Lower panel depicts recombination rate. **c** Colocalisation plot of the cg16885113 im-meQTL with lung cancer. **d** Correlation of rs6446553 genotype with *ZFP57* expression in unstimulated monocytes (green) and following 2 and 24 hours of LPS stimulation (red). NS, not significant. **e** Regional association plot of *ZFP57* eQTL in untreated monocytes (n=414). Protein-coding genes are highlighted (blue). Lower panel depicts recombination rate. **f** Circos plot highlighted CpGs with methylation levels significantly modified by *ZFP57* eQTL genotype. Known imprinted regions are high-lighted (orange). **g** Correlation of baseline CpG methylation levels and the effect of *ZFP57* eQTL genotype (rs417764) on methylation at CpGs significantly affected by rs417764 genotype (n=151, *r* = *−*0.33, *p* = 3.0 *×* 10*^−^*^5^). **h** Distribution of methylation levels in untreated monocytes at *ZFP57* eQTL-associated CpGs according to genotype at rs417764 (n=151). **i** Correlation of *ZFP57* eQTL genotype (rs417764) with change in methylation at significantly rs417764-associated CpGs (n=7). For regional association and colocalisation plots, SNPs are coloured according to strength of LD (CEU population) to the peak im-meQTL/eQTL SNP. Box and whisker plots; boxes depict the upper and lower quartiles of the data, and whiskers depict the range of the data excluding outliers (outliers are defined as data-points *>* 1.5*×* the inter-quartile range from the upper or lower quartiles).

Average levels of methylation at each CpG significantly modulated by rs417764 genotype are negatively correlated with the effect of rs417764:A allele carriage on CpG methylation (*r* = *−*0.33, *p* = 3.0 *×* 10*^−^*^5^; Fig. 5g). This effect is largely driven by genotype-dependent demethylation of highly methylated CpGs, which is associated with increased *ZFP57* expression (Fig 5g,h). Given that baseline methylation state is an important predictor of differential methylation in response to LPS stimulation, and that imCpGs are more likely to have intermediate methylation levels at baseline, we further hypothesised that the *ZFP57* eQTL would modify LPS responsiveness at CpGs where rs417764 was a predictor of baseline methylation. Of 151 ZFP57-modified CpGs, 7 have a significant effect (*FDR <* 0.05) of rs417764 genotype on differential methylation in response to LPS stimulation (Fig. 5i, Table S22). In keeping with the effect of rs417764 genotype on baseline methylation, where the rs417764:A allele is associated with more intermediate methylation levels at baseline, at 6 of these 7 CpGs the effect of rs417764:A allele carriage is to increase the magnitude of change in methylation in response to LPS stimulation (Fig. 5i).

## Discussion

In this study we explored the dynamics of DNAm in primary human monocytes following innate immune stimulation. Of the innate immune stimuli we tested, LPS induced the most marked changes in DNAm. This is in contrast to the effects of LPS and IFN*γ* on gene expression, where either stimulus induces changes of equivalent magnitude in the monocyte transcriptome[38]. We explored the DNAm response to LPS in a large cohort of healthy European-ancestry adults finding imCpGs were both methylated and demethylated in response to LPS, and LPS-induced demethylation was an active process, involving the formation of 5hmC. The sites at which LPS-induced methylation and demethylation occurred were distinct: demethylated imCpGs demonstrating striking enrichment for enhancers and transcriptionally active DNA, whereas methylated imCpGs were found largely in transcriptionally silent regions. This observation lends itself to a hypothesis whereby LPS-induced demethylation in monocytes acts to further upregulate inflammatory pathways already active at baseline, whereas LPS-induced methylation further represses quiescent transcriptional programmes. In keeping with this, demethylated imCpGs are highly enriched for AP-1 subunit binding sites in myeloid cells, and are proximal to genes directing innate immune activity, including TLR signalling. By contrast, transcription factor usage at methylated imCpGs is modest and is dominated by under-representation at transcription factor binding sites, and consistent with this, genes proximal to these sites are not enriched for immunological signalling pathways. That is, transcriptional programmes modified by LPS-induced demethylation are specific, whereas those modified by LPS-induced methylation are more diverse and not representative of a cohesive set of biological processes.

We found that epigenetic age, as estimated by the Horvath DNAm clock[34], is accelerated by LPS stimulation in monocytes, and that this effect is only seen among individuals without baseline epigenetic age acceleration. This demonstrates that epigenetic age is modifiable in the short-term by innate immune stimulation, and suggests a model in which infectious and inflammatory exposures across the life-course contribute to epigenetic aging. Importantly, epigenetic age acceleration is associated with a variety of health outcomes, including cancer[11]. Complementary to this, LPS-demethylated imCpGs were found to be proximal to genes involved in human disease pathogenesis, in particular carcinogenesis. Moreover, the most markedly LPS-demethylated region is at *CUX1*, a gene recurrently mutated in leukaemias[22, 23]. The association between LPS-stimulation, accelerated epigenetic age and human disease speaks to the established association of chronic inflammation and age-related disease traits[1]. Cellular senescence contributes to the accumulation of systemic chronic inflammation with age[51], but chronic antigenic stimulation, in particular chronic viral infection, is also a key driver of immunosenescence[52] as well as acceleration of epigenetic age[53, 54]. However, the specific enrichment we observed for cancer risk suggests the possibility that LPS-induced demethylation plays a direct role in carcinogenesis. CpG dinucleotides accumulate mutation at rates greater than non-CpG DNA[55], and this interacts with methylation status[56]. It is plausible, therefore, that inflammation-induced methylation changes affect cancer risk through modified transcriptional programmes and cellular differentiation, but may also be directly mutagenic, potentially through occasional fallibility of demethylation associated base excision-repair.

Finally, we used eQTL mapping to define local genetic variation determining LPS-induced changes in DNAm at imCpGs. We were able to identify 209 instances of *cis* im-mQTL. In keeping with the observed enrichment of disease-relevant genes among genes proximal to imCpGs, im-mQTL colocalise with genetic determinants of human phenotypic variation and disease traits, again including cancer-associated loci. We further identified an im-mQTL which is also a determinant of *ZFP57* RNA expression in untreated monocytes. ZFP57 controls bistability at imprinted loci[43–45], and in keeping with this we were able to observe effects of the *ZFP57* eQTL on baseline methlyation at 151 CpG dinucleotides genome-wide, which were highly enriched for known imprinted loci. Consistent with the importance of baseline methylation in determining the DNAm LPS response, the *ZFP57* eQTL also predicts DNAm LPS response at a subset of these loci, demonstrating the intricate genome-wide relationship between genotype, gene expression, DNA methylation and environment.

In summary, we have described how the landscape of DNAm in primary human monocytes is modified by innate immune stimulation, demonstrating for the first time that these responses are in part genetically determined. In so doing, we have highlighted that monocyte DNAm and associated epigenetic age is modifiable in the short-term by acute inflammation, with changes being informative to the pathogenesis of a range of human disease states, most notably cancer.

## Methods

### Cell purification & stimulation

Peripheral blood mononuclear cells were separated from freshly drawn blood using Ficoll gradient purification, with monocytes subsequently positively selected using magnetic CD14^+^ isolation kits (miltenyi) according to manufacturer’s protocols. Purity of the monocytes was assessed from methylation values using the Houseman[57] method within the minfi[58] R package and was found to be at a median of *>* 99% for all treatments (Fig. S7). Monocytes were cultured at 500,000 cells per ml in 400*µ*l RPMI supplemented with L-Glutamine, Penicillin/Streptomycin and 20% FCS in BD Falcon 5ml polypropylene culture tubes (cat number 352063), with each sample typically being purified from 800,000 cells. Post purification samples were rested overnight at 37C, 5% CO 2 prior to stimulation under the same conditions with the following reagents and concentrations - Pam3Csk4 (Invivogen, Lot no.), LPS (Invivogen, Lot No.), IFN*γ* (). To determine whether stimulation elicited cell division, we used the CellTrace kit (Thermofisher) and subsequent flow cytometry according to manufacturer’s instructions in cells from four individuals.

### Array Methylation Analysis

DNA and RNA were purified from cells using the Qiagen kit () according to manufacturers instructions. DNA was bisulfite converted (Zymogen?) prior to hybridization to Illumina 450K arrays which carries 485,000 probes. For the LPS analysis, all but 6 treated samples were hybridized with their untreated control to an array on the same beadchip, thus mitigating chip and batch effects as far as possible. The Illumina 450K array carries a subset of SNPs and these were used to detect sample mix-ups. After quality control, we used a total of 380 paired samples from 190 individuals in the final analysis to assess the effect of 24 hours of LPS stimulation on DNA methylation across a population. Samples were normalized using quantile normalization using the EWAS pipeline described by Lehne, Drong *et al* [59]. CpGs with detection *p >* 1 *×* 10*^−^*^16^ in any sample were excluded from analysis. In addition, we excluded probes previously found to blat to more than one genomic location or overlapping SNPs with minor allele frequencies (MAF) *>* 1%[60]. This resulted in 407,951 probes and subsequent pairwise analysis was performed on normalized data using limma[15]. We identified differentially methylated regions using DMRcate[20] using default settings.

### Bisulfite sequencing and hydroxymethylation analysis

For four individuals, monocytes were treated for 0,6 or 24h of LPS prior to DNA purification (1 µg per sample) with subsequent paired bisulfite and oxidative bisulfite conversion using the CEGX oxidative-bisulfite sequencing kit. Converted DNA was then subjected to capture enrichment for 80.5Mb across regions known to be informative as to methylation status (SeqCap Epi Enrichment, Nimblegen). Multiplexed samples were subsequently sequenced at 100bp paired end reads using the Illumina HiSeq2500 platform using the rapid mode.

### Genotyping and imputation

Genome-wide genotypes were generated at in study participants using the UK Biobank Axiom array (Affymetrix). Samples were excluded from downstream analysis if participants were related (relatedness coefficient ¿0.05) or with outlying heterozygosity and call rate. SNPs with low call rates (*<* 0.98), evidence for departure from Hardy-Weinberg equilibrium (*p <* 0.00001) and MAF *<* 0.01 were excluded from further processing. Following sample and SNP QC, genotypes at 623,652 autosomal SNPs in 186 individuals were taken forward for genome-wide imputation. Imputation was performed using minimac4[61], following phasing with Eagle2[62], as implemented by the Michigan Imputation Server[63], using the Haplotype Reference Consortium r1.1[64] as a reference panel. Genome-wide imputation resulted in high confidence (*r*^2^ *>* 0.7, HWE *p >* 1 *×* 10*^−^*^10^) genotypes at 5,411,481 common (*maf >* 0.04), autosomal SNPs, which were used in downstream association analysis.

### QTL mapping

We used QTLtools[65] to map genetic determinants of methylation. We mapped correlation between CpG methylation and genotype in an additive linear model, including genetic variants in *cis* (within 100kb) to each CpG tested. QTLtools controls FDR at the level of each phenotype (CpG) by approximating a permutation test, and we used 10,000 permutations in our *cis* QTL mapping. We applied a second level of FDR control across all phenotypes tested with qvalue in R. We considered *FDR <* 0.05 to be significant. We sought to map genetic determinants of changes in methylation on LPS stimulation, which we term immune-modulated methylation quantitative trait loci (im-meQTL), correlating genotype with normalised change in CpG methylation, i.e. delta beta methylation, at each CpG for which there was evidence of differential methylation on LPS stimulation (n=20,858). In addition, we mapped the *cis* genetic determinants of CpG methylation in untreated and LPS-stimulated monocytes separately, mapping QTL at 407,951 CpGs passing QC across all samples. To minimise the effect of confounding variation, we included principal components (PC) of the phenotype matrix as covariates in the linear model, with the number of PC used in each analysis chosen to maximise QTL discovery. Inclusion of 29 PCs optimised im-meQTL discovery (Fig. S8), while 28 and 23 PCs optimised meQTL discovery in untreated and LPS-stimulated monocytes respectively (Fig. S9). PC analysis of genotyping data was not suggestive of confounding population structure, and comparison of study sample genetic PCs with those of 1000G project reference samples confirmed European ancestry (Fig. S10). We did not therefore include genetic PCs as covariates for eQTL mapping.

### Approaches to enrichment and colocalisation

To assess the distribution of imCpGs with respect to genomic features of interest, we used Roadmap Epigenomics Consortium[25] chromatin state annotations in primary moncytes defined by the ChromHMM[66] 15 state model, ENCODE Chip-seq data defining transcription factor binding sites in myeloid lineage cells (K562) and lymphoblastoid cell lines (LCLs - GM12878), and promoter capture HiC data across 17 cell types[26] defining enhancer activity. We assessed enrichment of imCpGs against a background of all CpGs tested (passing QC) for each feature of interest in XGR[29], using a two-tailed binomial test. Following QTL mapping, we tested for enrichment of im-meQTL in K562 transcription factor binding sites using a permutation-based approach implemented in QTLtools[65], calculating the frequency of observed overlap between a given transcription factor binding site and an im-meQTL peak SNP, comparing this to the the number of overlaps expected by chance (permuting phenotypes across all imCpGs tested).

We assessed enrichment of genes proximal to imCpGs (¡5kb, taking the single most proximal protein-coding gene) and genes correlated with principal components of LPS-induced differential methylation at imCpGs within biological pathways using the following ontologies; Reactome Immune System, Disease Ontology and Gene Ontology Biological Processes. We tested enrichment in XGR[29] using Fisher’s exact tests.

To test if im-meQTL are associated with phenotypic traits or risk of human disease we used a Bayesian approach implemented in Coloc v5.1.0.1 to assess the probability of a shared causal variant between the im-meQTL and trait-associated variation identified by GWAS. We downloaded casecontrol GWAS summary statistics (n=45, Table S23) from the MRC IEU OpenGWAS project[67] and UK Biobank GWAS summary statistics (url: http://www.nealelab.is/uk-biobank/) of peripheral blood traits (n=27, Table S23). We tested for evidence of colocalisation between each identified im-meQTL (n=209) and each GWAS trait within a 250kb window centred on the peak im-meQTL eSNP. To assess evidence for shared causal loci between im-meQTL and regulatory determinants of gene expression in untreated and LPS-stimulated monocytes, we used the multi-trait extension of Coloc, moloc v0.1.0[39]. We used previously-published summary statistics describing eQTL mapping of RNA expression in untreated monocytes (n=414), 2 hours LPS-treated monocytes (n=261) and 24 hours LPS treated monocytes (n=322)[38]. We tested for evidence of colocalisation between each identified im-meQTL and the 3 monocyte eQTL datasets within a 250kb window centred on the peak im-meQTL eSNP. We used default priors in all colocalisation analyses, and we considered a posterior probability supporting a shared causal locus *>* 0.8 to be significant.

### Data and code availability

Raw methylation files will be made openly accessible via EGA. Genotyping data will be available upon completion of a data transfer agreement. Code to make figures will be made available on the group bitbucket https://bitbucket.org/Fairfaxlab/.

### Author Contributions

The study was conceived by BPF and JCK, who jointly oversaw the project. Samples were collected by SD, EL, BPF, HAM with access to the biobank provided by MN. Primary analysis was performed by JJG, BPF, MT, HF, IN, CT and OT. The manuscript was drafted by BPF and JJG with input and revisions from all other authors.

## Acknowledgments

The study was funded by Wellcome Trust Intermediate Clinical Fellowship to B.P.F. (no. 201488/Z/16/Z). J.C.K. is supported by Wellcome Trust Investigator Award [204969/Z/16/Z], NIHR Oxford Biomedical Research Centre and Chinese Academy of Medical Sciences (CAMS) Innovation Fund for Medical Science (grant number: 2018-I2M-2-002), Wellcome Trust Grants 090532/Z/09/Z and 203141/Z/16/Z to core facilities Wellcome Centre for Human Genetics, Oxford Biomedical Research Computing (BMRC) facility, a joint development between the Wellcome Centre for Human Genetics and the Big Data Institute supported by Health Data Research UK and the NIHR Oxford Biomedical Research Centre. J.J.G. is funded by a National Institute for Health Research (NIHR) Clinical Lectureship. The views expressed are those of the author(s) and not necessarily those of the NHS, the NIHR or the Department of Health and Social Care.

---

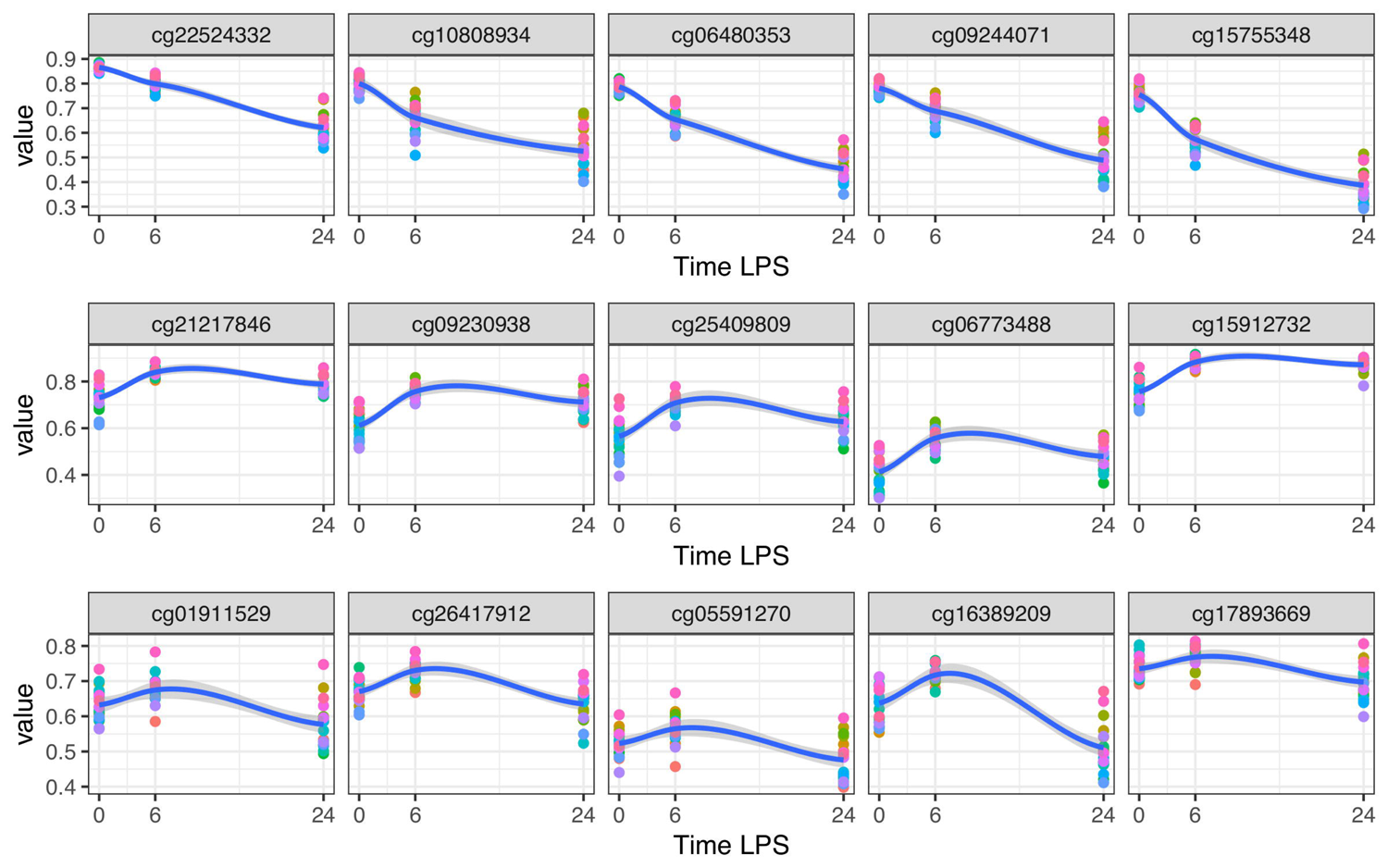

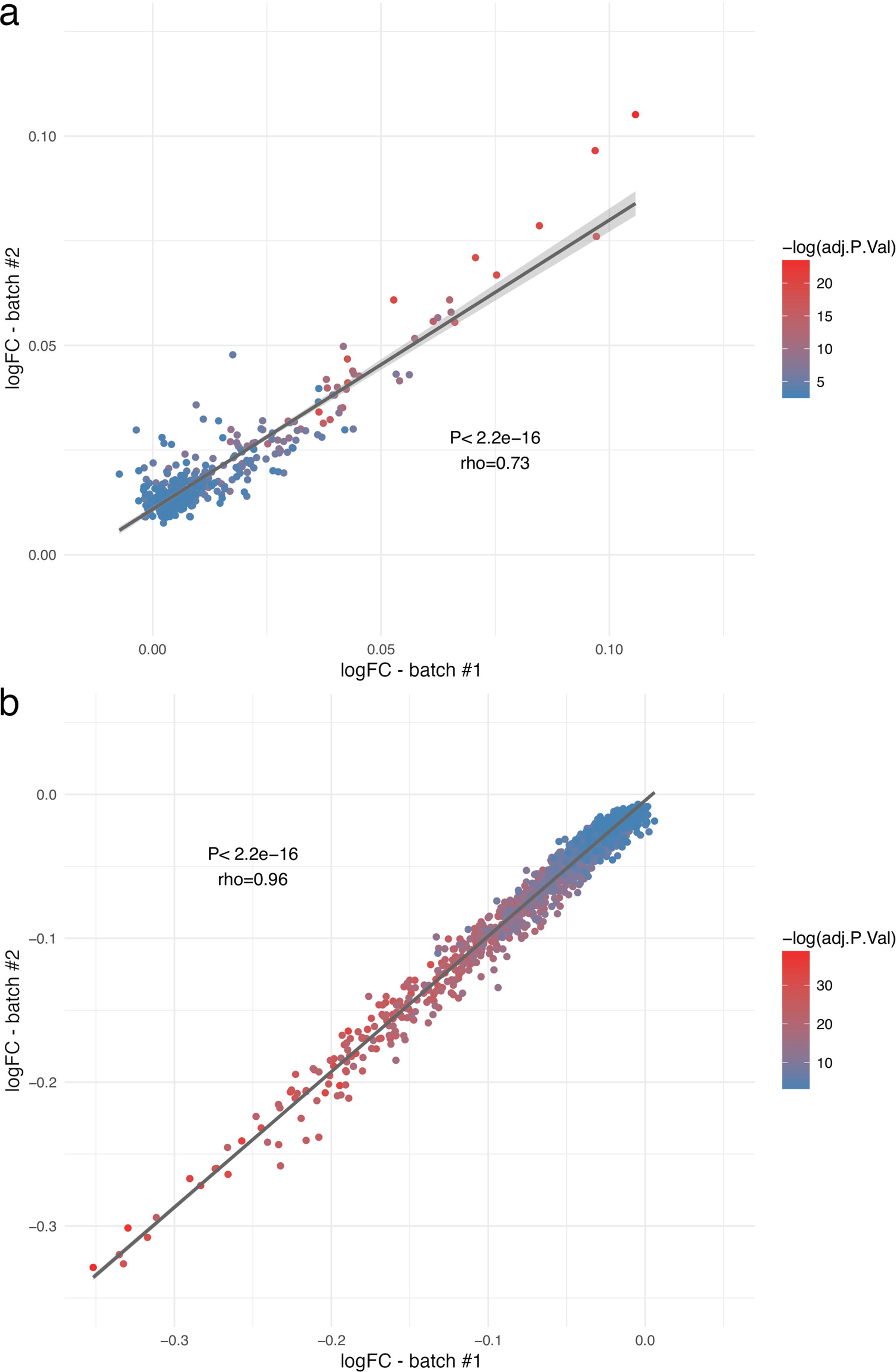

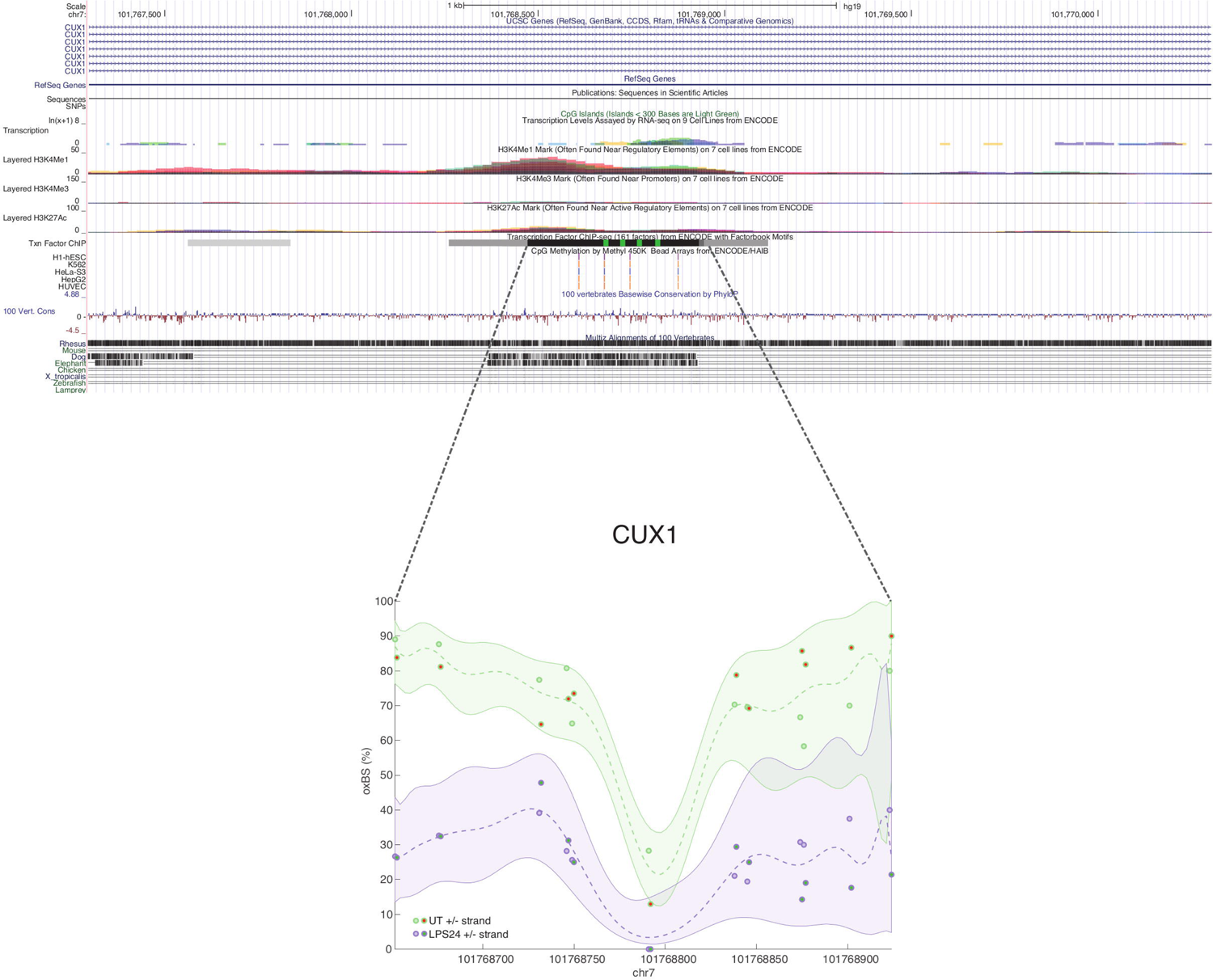

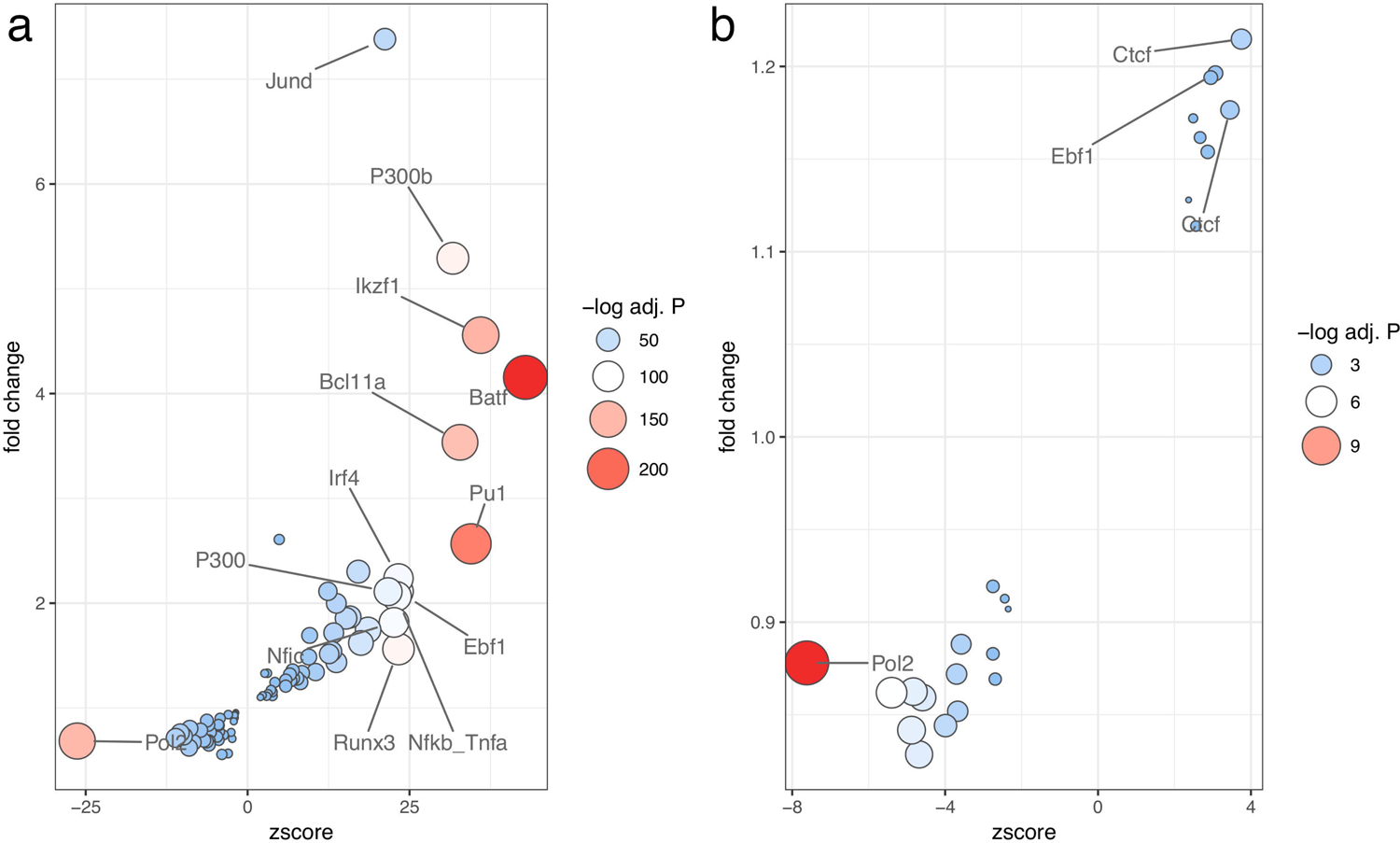

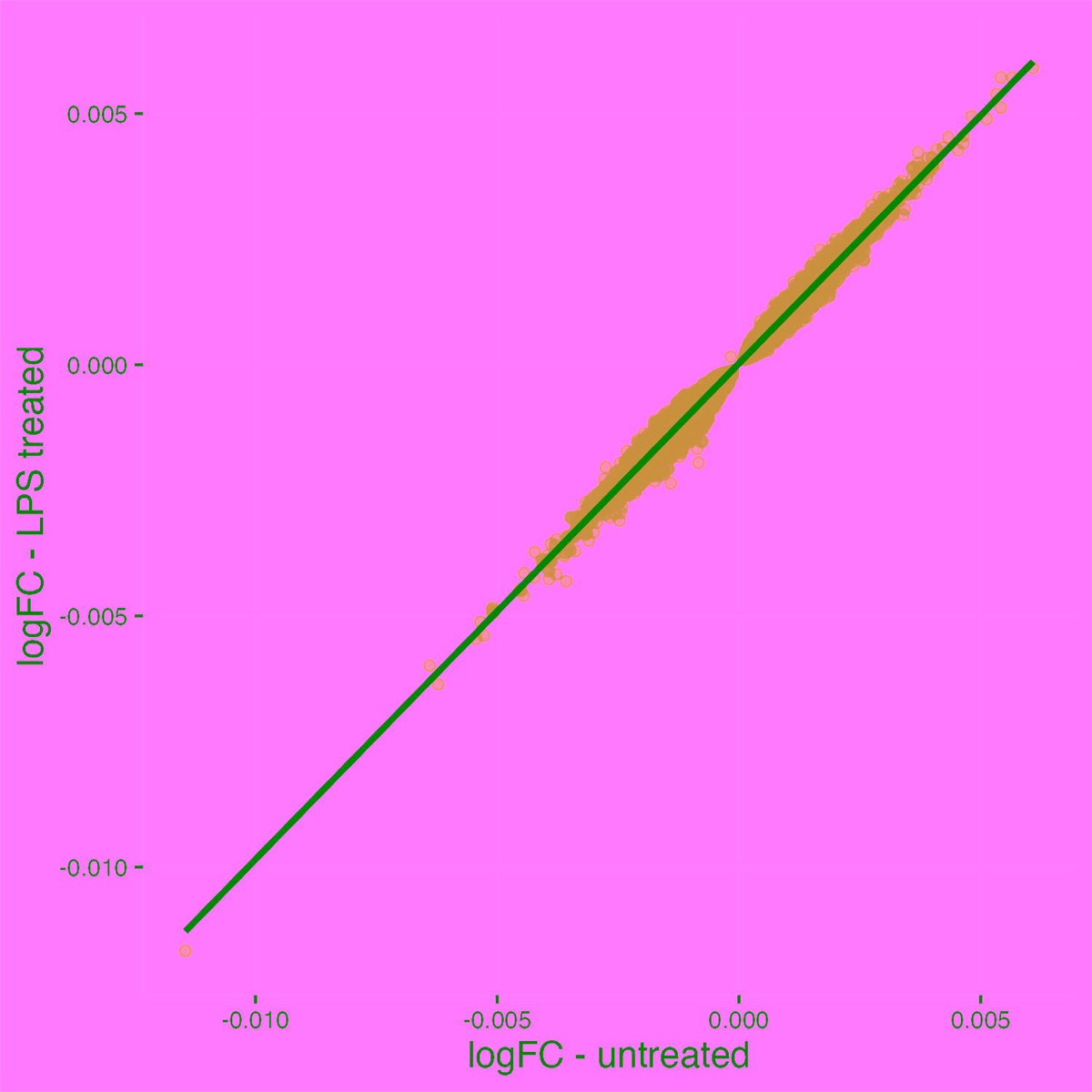

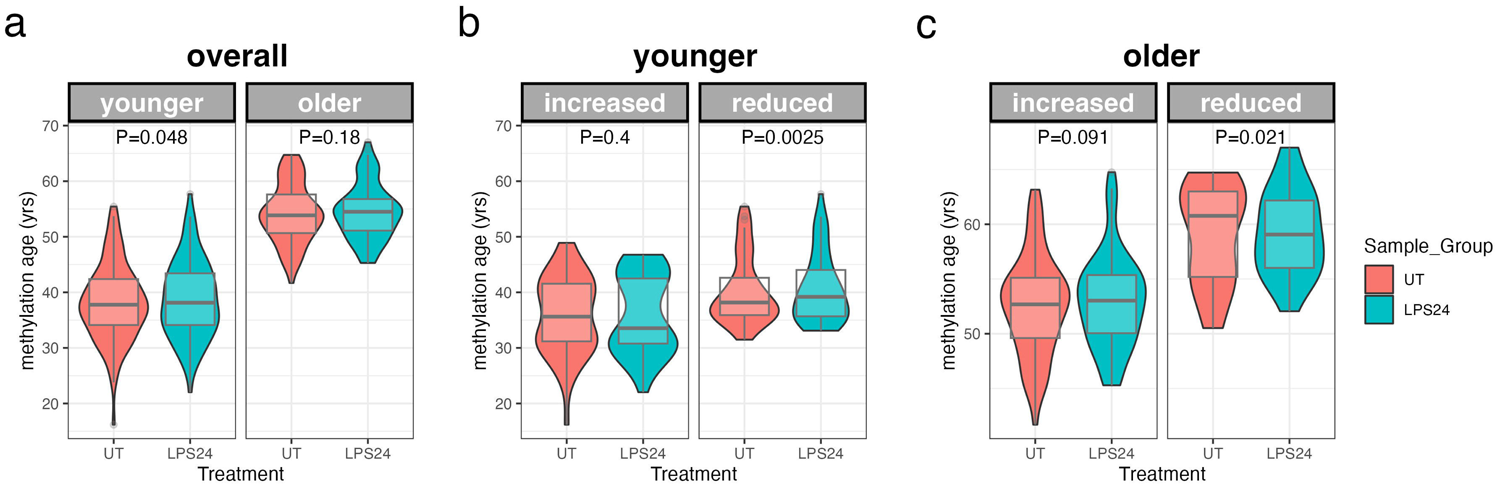

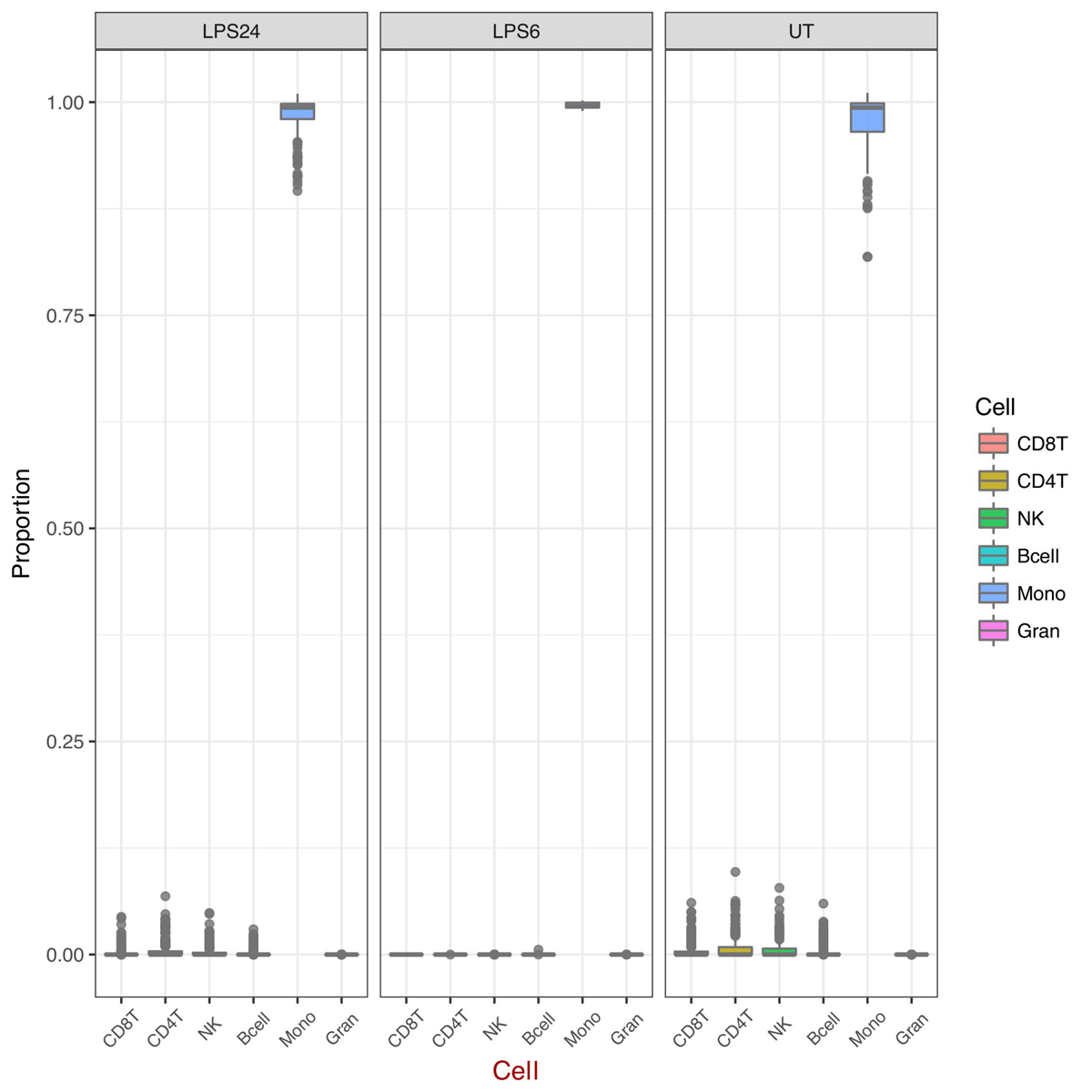

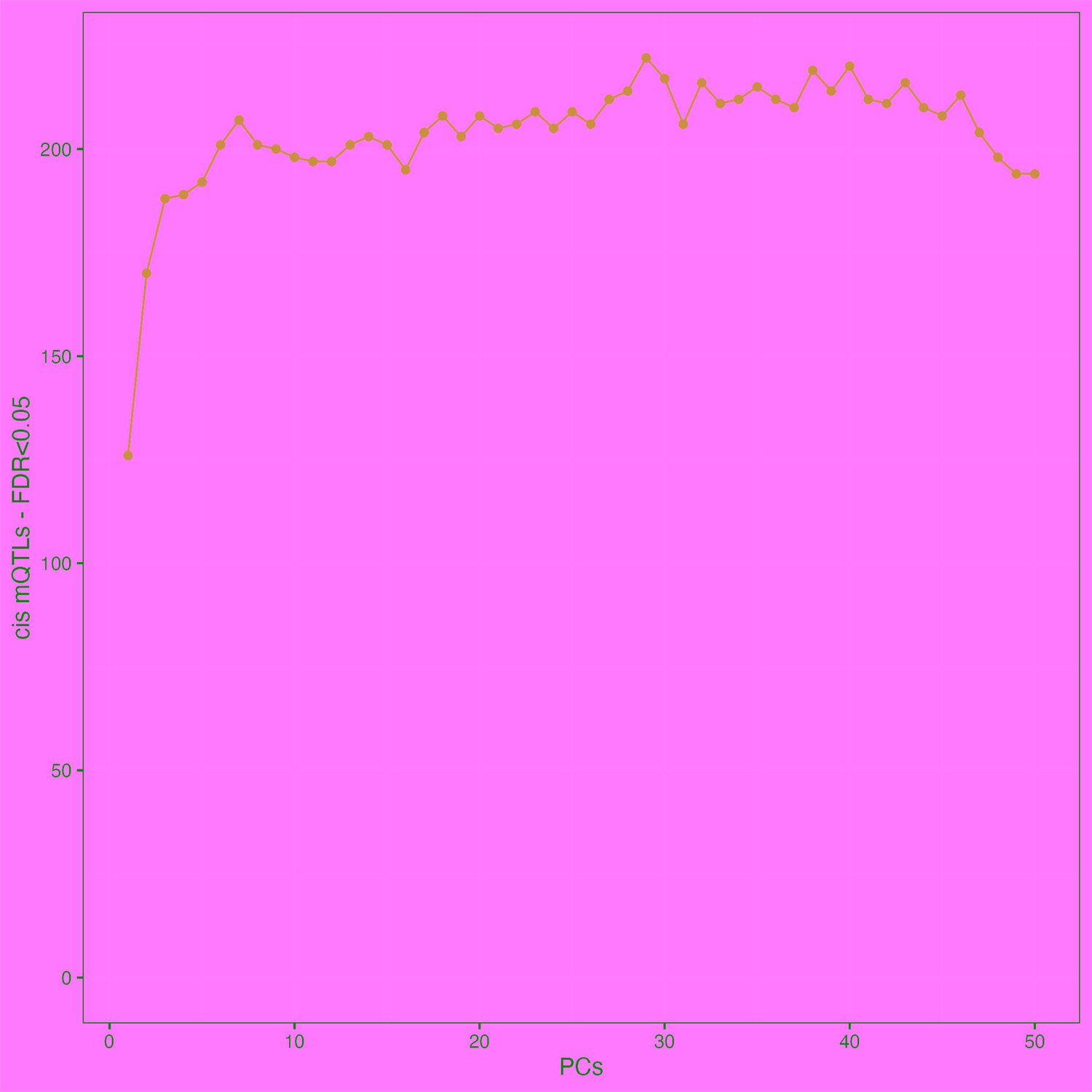

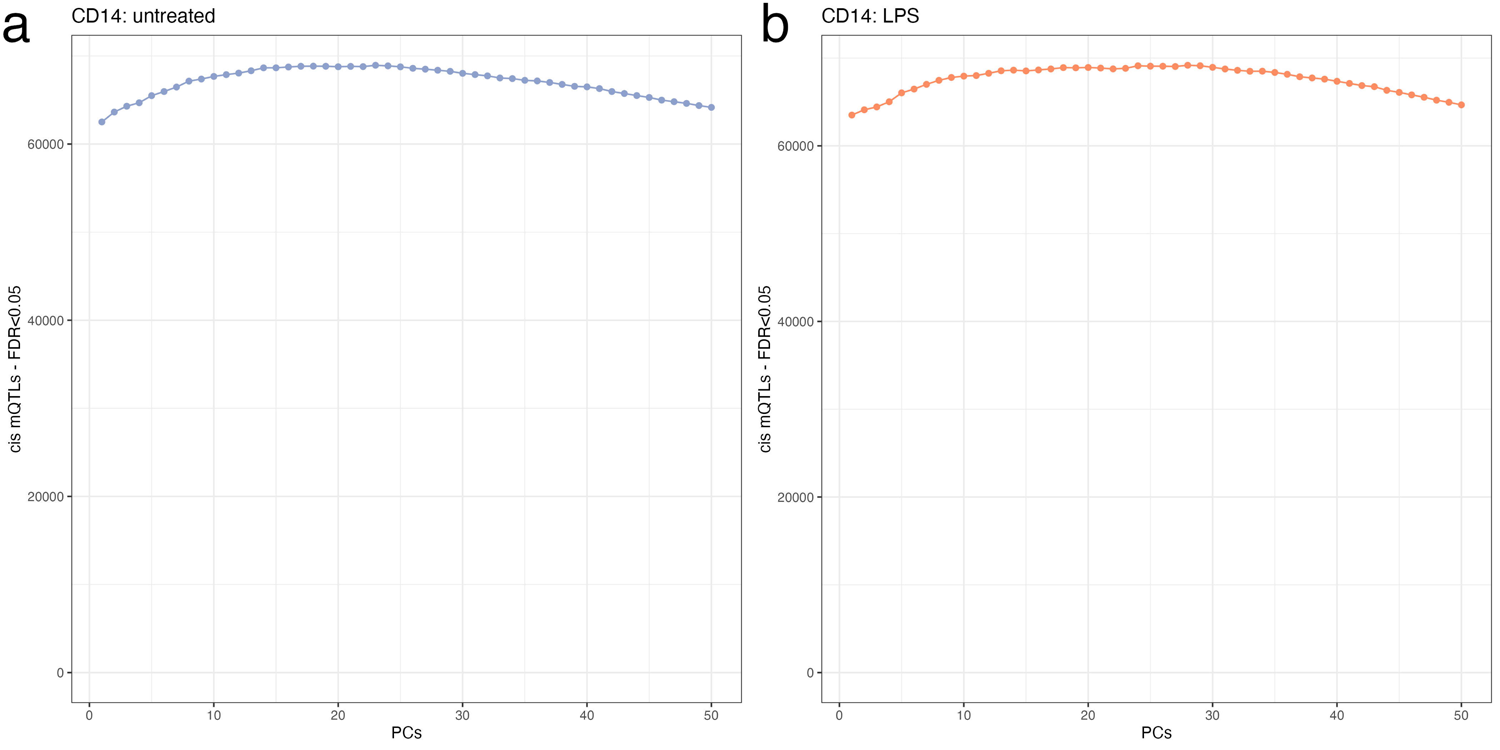

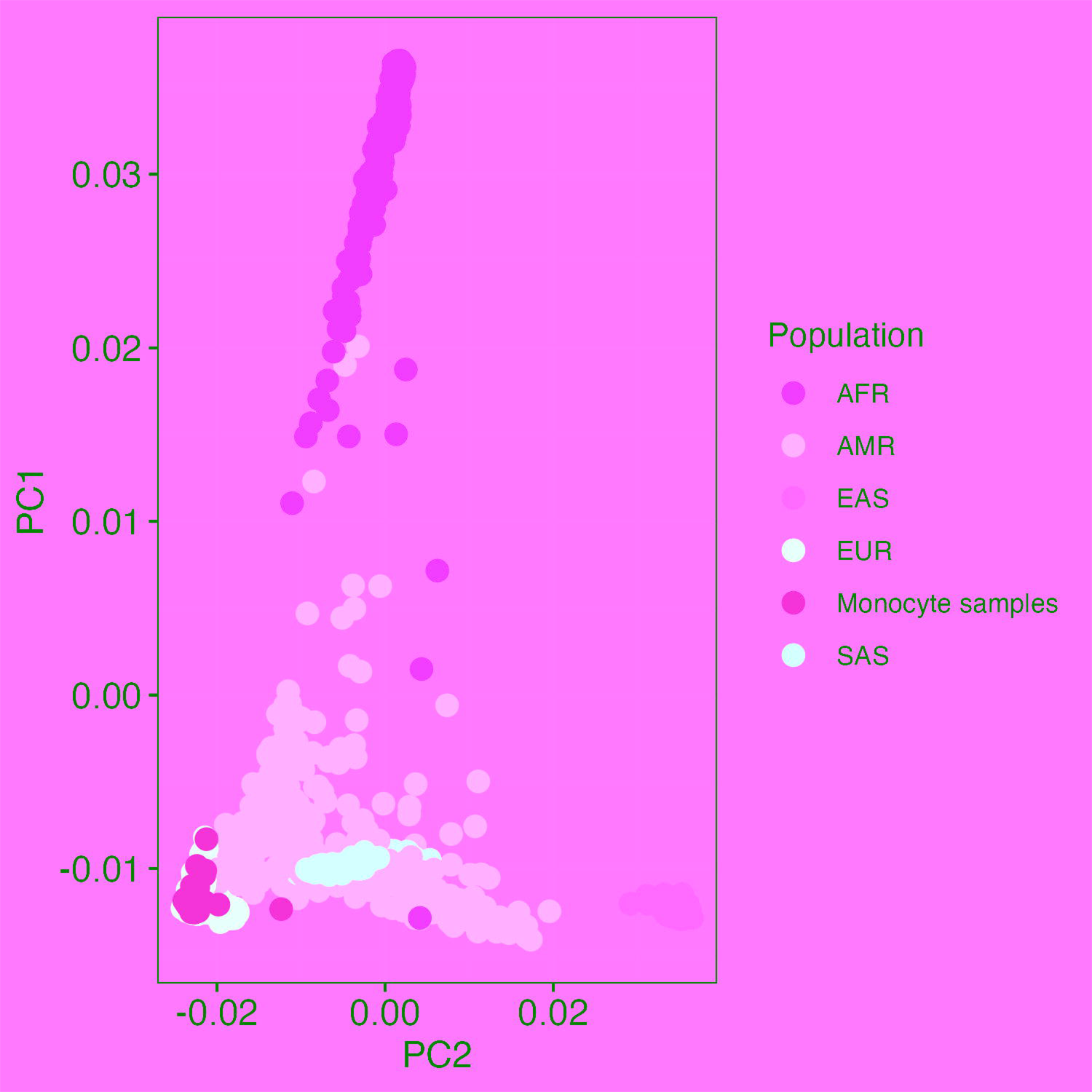

## References

[1] Furman, D., Campisi, J., Verdin, E., Carrera-Bastos, P., Targ, S., Franceschi, C., Ferrucci, L., Gilroy, D.W., Fasano, A., Miller, G.W., Miller, A.H., Mantovani, A., Weyand, C.M., Barzilai, N., Goronzy, J.J., Rando, T.A., Effros, R.B., Lucia, A., Kleinstreuer, N., Slavich, G.M.: Chronic inflammation in the etiology of disease across the life span. Nat Med 25(12), 1822–1832 (2019). https://doi.org/10.1038/s41591-019-0675-0

[2] Schübeler, D.: Function and information content of dna methylation. Nature 517(7534), 321–326 (2015). https://doi.org/10.1038/nature14192

[3] Grundberg, E., Meduri, E., Sandling, J.K., Hedman, A.K., Keildson, S., Buil, A., Busche, S., Yuan, W., Nisbet, J., Sekowska, M., Wilk, A., Barrett, A., Small, K.S., Ge, B., Caron, M., Shin, S.-Y., Lathrop, M., Dermitzakis, E.T., McCarthy, M.I., Spector, T.D., Bell, J.T., Deloukas, P.: Global analysis of dna methylation variation in adipose tissue from twins reveals links to disease-associated variants in distal regulatory elements. Am J Hum Genet 93(5), 876–890 (2013). https://doi.org/10.1016/j.ajhg.2013.10.004

[4] Bonder, M.J., Luijk, R., Zhernakova, D.V., Moed, M., Deelen, P., Vermaat, M., van Iterson, M., van Dijk, F., van Galen, M., Bot, J., Slieker, R.C., Jhamai, P.M., Verbiest, M., Suchiman, H.E.D., Verkerk, M., van der Breggen, R., van Rooij, J., Lakenberg, N., Arindrarto, W., Kielbasa, S.M., Jonkers, I., van’t Hof, P., Nooren, I., Beekman, M., Deelen, J., van Heemst, D., Zhernakova, A., Tigchelaar, E.F., Swertz, M.A., Hofman, A., Uitterlinden, A., Pool, R., van Dongen, J., Hottenga, J.J., Stehouwer, C.D.A., van der Kallen, C.J.H., Schalkwijk, C.G., van den Berg, L.H., van Zwet, E.W., Mei, H., Li, Y., Lemire, M., Hudson, T.J., Slagboom, P.E., Wijmenga, C., Veldink, J.H., van Greevenbroek, M.M.J., van Duijn, C.M., Boomsma, D.I., Isaacs, A., Jansen, R., van Meurs, J.B.J., ’t Hoen, P.A.C., Franke, L., Heijmans, B.T.: Disease variants alter transcription factor levels and methylation of their binding sites. Nat Genet 49(1), 131–138 (2017). https://doi.org/10.1038/ng.3721

[5] Kulis, M., Merkel, A., Heath, S., Queiŕos, A.C., Schuyler, R.P., Castellano, G., Beekman, R., Raineri, E., Esteve, A., Clot, G., Verdaguer-Dot, N., Duran-Ferrer, M., Russiñol, N., Vilarrasa-Blasi, R., Ecker, S., Pancaldi, V., Rico, D., Agueda, L., Blanc, J., Richardson, D., Clarke, L., Datta, A., Pascual, M., Agirre, X., Prosper, F., Alignani, D., Paiva, B., Caron, G., Fest, T., Muench, M.O., Fomin, M.E., Lee, S.-T., Wiemels, J.L., Valencia, A., Gut, M., Flicek, P., Stunnenberg, H.G., Siebert, R., Küppers, R., Gut, I.G., Campo, E., Martín-Subero, J.: Whole-genome fingerprint of the dna methylome during human b cell differentiation. Nat Genet 47(7), 746–756 (2015). https://doi.org/10.1038/ng.3291

[6] Ladle, B.H., Li, K.-P., Phillips, M.J., Pucsek, A.B., Haile, A., Powell, J.D., Jaffee, E.M., Hildeman, D.A., Gamper, C.J.: De novo dna methylation by dna methyltransferase 3a controls early effector cd8+ t-cell fate decisions following activation. Proc Natl Acad Sci U S A 113(38), 10631–10636 (2016). https://doi.org/10.1073/pnas.1524490113

[7] Breitling, L.P., Yang, R., Korn, B., Burwinkel, B., Brenner, H.: Tobacco-smoking-related differential dna methylation: 27k discovery and replication. Am J Hum Genet 88(4), 450–457 (2011). https://doi.org/10.1016/j.ajhg.2011.03.003

[8] Horvath, S., Raj, K.: Dna methylation-based biomarkers and the epigenetic clock theory of ageing. Nat Rev Genet 19(6), 371–384 (2018). https://doi.org/10.1038/s41576-018-0004-3

[9] Horvath, S., Pirazzini, C., Bacalini, M.G., Gentilini, D., Di Blasio, A.M., Delledonne, M., Mari, D., Arosio, B., Monti, D., Passarino, G., De Rango, F., D’Aquila, P., Giuliani, C., Marasco, E., Collino, S., Descombes, P., Garagnani, P., Franceschi, C.: Decreased epigenetic age of pbmcs from italian semi-supercentenarians and their offspring. Aging (Albany NY) 7(12), 1159–1170 (2015). https://doi.org/10.18632/aging.100861

[10] Marioni, R.E., Shah, S., McRae, A.F., Ritchie, S.J., Muniz-Terrera, G., Harris, S.E., Gibson, J., Redmond, P., Cox, S.R., Pattie, A., Corley, J., Taylor, A., Murphy, L., Starr, J.M., Horvath, S., Visscher, P.M., Wray, N.R., Deary, I.J.: The epigenetic clock is correlated with physical and cognitive fitness in the lothian birth cohort 1936. Int J Epidemiol 44(4), 1388–1396 (2015). https://doi.org/10.1093/ije/dyu277

[11] Zheng, Y., Joyce, B.T., Colicino, E., Liu, L., Zhang, W., Dai, Q., Shrubsole, M.J., Kibbe, W.A., Gao, T., Zhang, Z., Jafari, N., Vokonas, P., Schwartz, J., Baccarelli, A.A., Hou, L.: Blood epigenetic age may predict cancer incidence and mortality. EBioMedicine 5, 68–73 (2016). https://doi.org/10.1016/j.ebiom.2016.02.008

[12] Marioni, R.E., Shah, S., McRae, A.F., Chen, B.H., Colicino, E., Harris, S.E., Gibson, J., Henders, A.K., Redmond, P., Cox, S.R., Pattie, A., Corley, J., Murphy, L., Martin, N.G., Montgomery, G.W., Feinberg, A.P., Fallin, M.D., Multhaup, M.L., Jaffe, A.E., Joehanes, R., Schwartz, J., Just, A.C., Lunetta, K.L., Murabito, J.M., Starr, J.M., Horvath, S., Baccarelli, A.A., Levy, D., Visscher, P.M., Wray, N.R., Deary, I.J.: Dna methylation age of blood predicts all-cause mortality in later life. Genome Biol 16(1), 25 (2015). https://doi.org/10.1186/s13059-015-0584-6

[13] Marr, A.K., MacIsaac, J.L., Jiang, R., Airo, A.M., Kobor, M.S., McMaster, W.R.: Leishmania donovani infection causes distinct epigenetic dna methylation changes in host macrophages. PLoS Pathog 10(10), 1004419 (2014). https://doi.org/10.1371/journal.ppat.1004419

[14] Pacis, A., Tailleux, L., Morin, A.M., Lambourne, J., MacIsaac, J.L., Yotova, V., Dumaine, A., Danckaert, A., Luca, F., Grenier, J.-C., Hansen, K.D., Gicquel, B., Yu, M., Pai, A., He, C., Tung, J., Pastinen, T., Kobor, M.S., Pique-Regi, R., Gilad, Y., Barreiro, L.B.: Bacterial infection remodels the dna methylation landscape of human dendritic cells. Genome Res 25(12), 1801–1811 (2015). https://doi.org/10.1101/gr.192005.115

[15] Ritchie, M.E., Phipson, B., Wu, D., Hu, Y., Law, C.W., Shi, W., Smyth, G.K.: limma powers differential expression analyses for rna-sequencing and microarray studies. Nucleic Acids Res 43(7), 47 (2015). https://doi.org/10.1093/nar/gkv007

[16] Ecker, S., Chen, L., Pancaldi, V., Bagger, F.O., Fernández, J.M., Carrillo de Santa Pau, E., Juan, D., Mann, A.L., Watt, S., Casale, F.P., et al.: Genome-wide analysis of differential transcriptional and epigenetic variability across human immune cell types. Genome biology 18, 1–17 (2017)

[17] Kohli, R.M., Zhang, Y.: Tet enzymes, tdg and the dynamics of dna demethylation. Nature 502(7472), 472–479 (2013). https://doi.org/10.1038/nature12750

[18] Tahiliani, M., Koh, K.P., Shen, Y., Pastor, W.A., Bandukwala, H., Brudno, Y., Agarwal, S., Iyer, L.M., Liu, D.R., Aravind, L., Rao, A.: Conversion of 5-methylcytosine to 5-hydroxymethylcytosine in mammalian dna by mll partner tet1. Science 324(5929), 930–935 (2009). https://doi.org/10.1126/science.1170116

[19] Booth, M.J., Ost, T.W.B., Beraldi, D., Bell, N.M., Branco, M.R., Reik, W., Balasubramanian, S.: Oxidative bisulfite sequencing of 5-methylcytosine and 5-hydroxymethylcytosine. Nat Protoc 8(10), 1841– 1851 (2013). https://doi.org/10.1038/nprot.2013.115

[20] Peters, T.J., Buckley, M.J., Statham, A.L., Pidsley, R., Samaras, K., V Lord, R., Clark, S.J., Molloy, P.L.: De novo identification of differentially methylated regions in the human genome. Epigenetics Chromatin 8, 6 (2015). https://doi.org/10.1186/1756-8935-8-6

[21] Vadnais, C., Awan, A.A., Harada, R., Clermont, P.-L., Leduy, L., Bérubé, G., Nepveu, A.: Long-range transcriptional regulation by the p110 cux1 homeodomain protein on the encode array. BMC Genomics 14, 258 (2013). https://doi.org/10.1186/1471-2164-14-258

[22] McNerney, M.E., Brown, C.D., Wang, X., Bartom, E.T., Karmakar, S., Bandlamudi, C., Yu, S., Ko, J., Sandall, B.P., Stricker, T., Anastasi, J., Grossman, R.L., Cunningham, J.M., Le Beau, M.M., White, K.P.: Cux1 is a haploinsufficient tumor suppressor gene on chromosome 7 frequently inactivated in acute myeloid leukemia. Blood 121(6), 975–983 (2013). https://doi.org/10.1182/blood-2012-04-426965

[23] Wong, C.C., Martincorena, I., Rust, A.G., Rashid, M., Alifrangis, C., Alexandrov, L.B., Tiffen, J.C., Kober, C., Green, A.R., Massie, C.E., Nangalia, J., Lempidaki, S., Döhner, H., Döhner, K., Bray, S.J., McDermott, U., Papaemmanuil, E., Campbell, P.J., Adamss, D.J.: Inactivating cux1 mutations promote tumorigenesis. Nat Genet 46(1), 33–38 (2014). https://doi.org/10.1038/ng.2846

[24] Al-Mossawi, H., Yager, N., Taylor, C.A., Lau, E., Danielli, S., de Wit, J., Gilchrist, N.I. James, Mahe, E.A., Lee, W., Rizvi, L., et al.: Context-specific regulation of surface and soluble il7r expression by an autoimmune risk allele. Nature Communications 10(4575) (2019)

[25] Kundaje, A., Meuleman, W., Ernst, J., Bilenky, M., Yen, A., Heravi-Moussavi, A., Kheradpour, P., Zhang, Z., Wang, J., Ziller, M.J., Amin, V., Whitaker, J.W., Schultz, M.D., Ward, L.D., Sarkar, A., Quon, G., Sandstrom, R.S., Eaton, M.L., Wu, Y.-C., Pfenning, A.R., Wang, X., Claussnitzer, M., Liu, Y., Coarfa, C., Harris, R.A., Shoresh, N., Epstein, C.B., Gjoneska, E., Leung, D., Xie, W., Hawkins, R.D., Lister, R., Hong, C., Gascard, P., Mungall, A.J., Moore, R., Chuah, E., Tam, A., Canfield, T.K., Hansen, R.S., Kaul, R., Sabo, P.J., Bansal, M.S., Carles, A., Dixon, J.R., Farh, K.-H., Feizi, S., Karlic, R., Kim, A.-R., Kulkarni, A., Li, D., Lowdon, R., Elliott, G., Mercer, T.R., Neph, S.J., Onuchic, V., Polak, P., Rajagopal, N., Ray, P., Sallari, R.C., Siebenthall, K.T., Sinnott-Armstrong, N.A., Stevens, M., Thurman, R.E., Wu, J., Zhang, B., Zhou, X., Beaudet, A.E., Boyer, L.A., De Jager, P.L., Farnham, P.J., Fisher, S.J., Haussler, D., Jones, S.J.M., Li, W., Marra, M.A., McManus, M.T., Sunyaev, S., Thomson, J.A., Tlsty, T.D., Tsai, L.-H., Wang, W., Waterland, R.A., Zhang, M.Q., Chadwick, L.H., Bernstein, B.E., Costello, J.F., Ecker, J.R., Hirst, M., Meissner, A., Milosavljevic, A., Ren, B., Stamatoyannopoulos, J.A., Wang, T., Kellis, M.: Integrative analysis of 111 reference human epigenomes. Nature 518(7539), 317–330 (2015). https://doi.org/10.1038/nature14248

[26] Javierre, B.M., Burren, O.S., Wilder, S.P., Kreuzhuber, R., Hill, S.M., Sewitz, S., Cairns, J., Wingett, S.W., Várnai, C., Thiecke, M.J., Burden, F., Farrow, S., Cutler, A.J., Rehnström, K., Downes, K., Grassi, L., Kostadima, M., Freire-Pritchett, P., Wang, F., Stunnenberg, H.G., Todd, J.A., Zerbino, D.R., Stegle, O., Ouwehand, W.H., Frontini, M., Wallace, C., Spivakov, M., Fraser, P.: Lineage-specific genome architecture links enhancers and non-coding disease variants to target gene promoters. Cell 167(5), 1369–1384 (2016). https://doi.org/10.1016/j.cell.2016.09.037

[27] Gillespie, M., Jassal, B., Stephan, R., Milacic, M., Rothfels, K., Senff-Ribeiro, A., Griss, J., Sevilla, C., Matthews, L., Gong, C., Deng, C., Varusai, T., Ragueneau, E., Haider, Y., May, B., Shamovsky, V., Weiser, J., Brunson, T., Sanati, N., Beckman, L., Shao, X., Fabregat, A., Sidiropoulos, K., Murillo, J., Viteri, G., Cook, J., Shorser, S., Bader, G., Demir, E., Sander, C., Haw, R., Wu, G., Stein, L., Hermjakob, H., D’Eustachio, P.: The reactome pathway knowledgebase 2022. Nucleic Acids Res 50(D1), 687–692 (2022). https://doi.org/10.1093/nar/gkab1028

[28] Schriml, L.M., Mitraka, E., Munro, J., Tauber, B., Schor, M., Nickle, L., Felix, V., Jeng, L., Bearer, C., Lichenstein, R., Bisordi, K., Campion, N., Hyman, B., Kurland, D., Oates, C.P., Kibbey, S., Sreekumar, P., Le, C., Giglio, M., Greene, C.: Human disease ontology 2018 update: classification, content and workflow expansion. Nucleic Acids Res 47(D1), 955–962 (2019). https://doi.org/10.1093/nar/gky1032

[29] Fang, H., Knezevic, B., Burnham, K.L., Knight, J.C.: Xgr software for enhanced interpretation of genomic summary data, illustrated by application to immunological traits. Genome Med 8(1), 129 (2016). https://doi.org/10.1186/s13073-016-0384-y

[30] Fraga, M.F., Ballestar, E., Paz, M.F., Ropero, S., Setien, F., Ballestar, M.L., Heine-Suñer, D., Cigudosa, J.C., Urioste, M., Benitez, J., Boix-Chornet, M., Sanchez-Aguilera, A., Ling, C., Carlsson, E., Poulsen, P., Vaag, A., Stephan, Z., Spector, T.D., Wu, Y.-Z., Plass, C., Esteller, M.: Epigenetic differences arise during the lifetime of monozygotic twins. Proc Natl Acad Sci U S A 102(30), 10604–10609 (2005). https://doi.org/10.1073/pnas.0500398102

[31] Rakyan, V.K., Down, T.A., Maslau, S., Andrew, T., Yang, T.-P., Beyan, H., Whittaker, P., McCann, O.T., Finer, S., Valdes, A.M., Leslie, R.D., Deloukas, P., Spector, T.D.: Human aging-associated dna hypermethylation occurs preferentially at bivalent chromatin domains. Genome Res 20(4), 434–439 (2010). https://doi.org/10.1101/gr.103101.109

[32] Teschendorff, A.E., Menon, U., Gentry-Maharaj, A., Ramus, S.J., Weisenberger, D.J., Shen, H., Campan, M., Noushmehr, H., Bell, C.G., Maxwell, A.P., Savage, D.A., Mueller-Holzner, E., Marth, C., Kocjan, G., Gayther, S.A., Jones, A., Beck, S., Wagner, W., Laird, P.W., Jacobs, I.J., Widschwendter, M.: Age-dependent dna methylation of genes that are suppressed in stem cells is a hallmark of cancer. Genome Res 20(4), 440–446 (2010). https://doi.org/10.1101/gr.103606.109

[33] Levine, M.E., Lu, A.T., Quach, A., Chen, B.H., Assimes, T.L., Bandinelli, S., Hou, L., Baccarelli, A.A., Stewart, J.D., Li, Y., Whitsel, E.A., Wilson, J.G., Reiner, A.P., Aviv, A., Lohman, K., Liu, Y., Ferrucci, L., Horvath, S.: An epigenetic biomarker of aging for lifespan and healthspan. Aging (Albany NY) 10(4), 573–591 (2018). https://doi.org/10.18632/aging.101414

[34] Horvath, S.: Dna methylation age of human tissues and cell types. Genome Biol 14(10), 115 (2013). https://doi.org/10.1186/gb-2013-14-10-r115

[35] Gopalan, S., Carja, O., Fagny, M., Patin, E., Myrick, J.W., McEwen, L.M., Mah, S.M., Kobor, M.S., Froment, A., Feldman, M.W., Quintana-Murci, L., Henn, B.M.: Trends in dna methylation with age replicate across diverse human populations. Genetics 206(3), 1659–1674 (2017). https://doi.org/10.1534/genetics.116.195594

[36] Sun, Z., Xu, X., He, J., Murray, A., Sun, M.-A., Wei, X., Wang, X., McCoig, E., Xie, E., Jiang, X., Li, L., Zhu, J., Chen, J., Morozov, A., Pickrell, A.M., Theus, M.H., Xie, H.: Egr1 recruits tet1 to shape the brain methylome during development and upon neuronal activity. Nat Commun 10(1), 3892 (2019). https://doi.org/10.1038/s41467-019-11905-3

[37] Gutierrez-Arcelus, M., Lappalainen, T., Montgomery, S.B., Buil, A., Ongen, H., Yurovsky, A., Bryois, J., Giger, T., Romano, L., Planchon, A., Falconnet, E., Bielser, D., Gagnebin, M., Padioleau, I., Borel, C., Letourneau, A., Makrythanasis, P., Guipponi, M., Gehrig, C., Antonarakis, S.E., Dermitzakis, E.T.: Passive and active dna methylation and the interplay with genetic variation in gene regulation. Elife 2, 00523 (2013). https://doi.org/10.7554/eLife.00523

[38] Fairfax, B.P., Humburg, P., Makino, S., Naranbhai, V., Wong, D., Lau, E., Jostins, L., Plant, K., Andrews, R., McGee, C., Knight, J.C.: Innate immune activity conditions the effect of regulatory variants upon mono-cyte gene expression. Science 343(6175), 1246949 (2014). https://doi.org/10.1126/science.1246949

[39] Giambartolomei, C., Zhenli Liu, J., Zhang, W., Hauberg, M., Shi, H., Boocock, J., Pickrell, J., Jaffe, A.E., Pasaniuc, B., Roussos, P.: A bayesian framework for multiple trait colocalization from summary association statistics. Bioinformatics 34(15), 2538–2545 (2018). https://doi.org/10.1093/bioinformatics/bty147

[40] Longatti, A., Lamb, C.A., Razi, M., Yoshimura, S.-i., Barr, F.A., Tooze, S.A.: Tbc1d14 regulates autophagosome formation via rab11- and ulk1-positive recycling endosomes. J Cell Biol 197(5), 659–675 (2012). https://doi.org/10.1083/jcb.201111079

[41] Ho, T.T., Warr, M.R., Adelman, E.R., Lansinger, O.M., Flach, J., Verovskaya, E.V., Figueroa, M.E., Passegúe, E.: Autophagy maintains the metabolism and function of young and old stem cells. Nature 543(7644), 205–210 (2017). https://doi.org/10.1038/nature21388

[42] Wang, Y., McKay, J.D., Rafnar, T., Wang, Z., Timofeeva, M.N., Broderick, P., Zong, X., Laplana, M., Wei, Y., Han, Y., Lloyd, A., Delahaye-Sourdeix, M., Chubb, D., Gaborieau, V., Wheeler, W., Chatterjee, N., Thorleifsson, G., Sulem, P., Liu, G., Kaaks, R., Henrion, M., Kinnersley, B., Valĺee, M., LeCalvez-Kelm, F., Stevens, V.L., Gapstur, S.M., Chen, W.V., Zaridze, D., Szeszenia-Dabrowska, N., Lissowska, J., Rudnai, P., Fabianova, E., Mates, D., Bencko, V., Foretova, L., Janout, V., Krokan, H.E., Gabrielsen, M.E., Skorpen, F., Vatten, L., Njølstad, I., Chen, C., Goodman, G., Benhamou, S., Vooder, T., Válk, K., Nelis, M., Metspalu, A., Lener, M., Lubiński, J., Johansson, M., Vineis, P., Agudo, A., Clavel-Chapelon, F., Bueno-de-Mesquita, H.B., Trichopoulos, D., Khaw, K.-T., Johansson, M., Weiderpass, E., Tjønneland, A., Riboli, E., Lathrop, M., Scelo, G., Albanes, D., Caporaso, N.E., Ye, Y., Gu, J., Wu, X., Spitz, M.R., Dienemann, H., Rosenberger, A., Su, L., Matakidou, A., Eisen, T., Stefansson, K., Risch, A., Chanock, S.J., Christiani, D.C., Hung, R.J., Brennan, P., Landi, M.T., Houlston, R.S., Amos, C.I.: Rare variants of large effect in brca2 and chek2 affect risk of lung cancer. Nat Genet 46(7), 736–741 (2014). https://doi.org/10.1038/ng.3002

[43] Butz, S., Schmolka, N., Karemaker, I.D., Villaseñor, R., Schwarz, I., Domcke, S., Uijttewaal, E.C.H., Jude, J., Lienert, F., Krebs, A.R., de Wagenaar, N.P., Bao, X., Zuber, J., Elling, U., Schübeler, D., Baubec, T.: Dna sequence and chromatin modifiers cooperate to confer epigenetic bistability at imprinting control regions. Nat Genet 54(11), 1702–1710 (2022). https://doi.org/10.1038/s41588-022-01210-z

[44] Mackay, D.J.G., Callaway, J.L.A., Marks, S.M., White, H.E., Acerini, C.L., Boonen, S.E., Dayanikli, P., Firth, H.V., Goodship, J.A., Haemers, A.P., Hahnemann, J.M.D., Kordonouri, O., Masoud, A.F., Oestergaard, E., Storr, J., Ellard, S., Hattersley, A.T., Robinson, D.O., Temple, I.K.: Hypomethylation of multiple imprinted loci in individuals with transient neonatal diabetes is associated with mutations in zfp57. Nat Genet 40(8), 949–951 (2008). https://doi.org/10.1038/ng.187

[45] Zuo, X., Sheng, J., Lau, H.-T., McDonald, C.M., Andrade, M., Cullen, D.E., Bell, F.T., Iacovino, M., Kyba, M., Xu, G., Li, X.: Zinc finger protein zfp57 requires its co-factor to recruit dna methyltransferases and maintains dna methylation imprint in embryonic stem cells via its transcriptional repression domain. J Biol Chem 287(3), 2107–2118 (2012). https://doi.org/10.1074/jbc.M111.322644

[46] Akbari, V., Garant, J.-M., O’Neill, K., Pandoh, P., Moore, R., Marra, M.A., Hirst, M., Jones, S.J.M.: Genome-wide detection of imprinted differentially methylated regions using nanopore sequencing. Elife 11 (2022). https://doi.org/10.7554/eLife.77898

[47] Court, F., Tayama, C., Romanelli, V., Martin-Trujillo, A., Iglesias-Platas, I., Okamura, K., Sugahara, N., Simön, C., Moore, H., Harness, J.V., Keirstead, H., Sanchez-Mut, J.V., Kaneki, E., Lapunzina, P., Soejima, H., Wake, N., Esteller, M., Ogata, T., Hata, K., Nakabayashi, K., Monk, D.: Genome-wide parent-of-origin dna methylation analysis reveals the intricacies of human imprinting and suggests a germline methylation-independent mechanism of establishment. Genome Res 24(4), 554–569 (2014). https://doi.org/10.1101/gr.164913.113

[48] Hernandez Mora, J.R., Tayama, C., Śanchez-Delgado, M., Monteagudo-Śanchez, A., Hata, K., Ogata, T., Medrano, J., Poo-Llanillo, M.E., Simön, C., Moran, S., Esteller, M., Tenorio, J., Lapunzina, P., Kagami, M., Monk, D., Nakabayashi, K.: Characterization of parent-of-origin methylation using the illumina infinium methylationepic array platform. Epigenomics 10(7), 941–954 (2018). https://doi.org/10.2217/epi-2017-0172

[49] Joshi, R.S., Garg, P., Zaitlen, N., Lappalainen, T., Watson, C.T., Azam, N., Ho, D., Li, X., Antonarakis, S.E., Brunner, H.G., Buiting, K., Cheung, S.W., Coffee, B., Eggermann, T., Francis, D., Geraedts, J.P., Gimelli, G., Jacobson, S.G., Le Caignec, C., de Leeuw, N., Liehr, T., Mackay, D.J., Montgomery, S.B., Pagnamenta, A.T., Papenhausen, P., Robinson, D.O., Ruivenkamp, C., Schwartz, C., Steiner, B., Stevenson, D.A., Surti, U., Wassink, T., Sharp, A.J.: Dna methylation profiling of uniparental disomy subjects provides a map of parental epigenetic bias in the human genome. Am J Hum Genet 99(3), 555–566 (2016). https://doi.org/10.1016/j.ajhg.2016.06.032

[50] Zink, F., Magnusdottir, D.N., Magnusson, O.T., Walker, N.J., Morris, T.J., Sigurdsson, A., Halldorsson, G.H., Gudjonsson, S.A., Melsted, P., Ingimundardottir, H., Kristmundsdottir, S., Alexandersson, K.F., Helgadottir, A., Gudmundsson, J., Rafnar, T., Jonsdottir, I., Holm, H., Eyjolfsson, G.I., Sigurdardottir, O., Olafsson, I., Masson, G., Gudbjartsson, D.F., Thorsteinsdottir, U., Halldorsson, B.V., Stacey, S.N., Stefansson, K.: Insights into imprinting from parent-of-origin phased methylomes and transcriptomes. Nat Genet 50(11), 1542–1552 (2018). https://doi.org/10.1038/s41588-018-0232-7

[51] Xu, M., Pirtskhalava, T., Farr, J.N., Weigand, B.M., Palmer, A.K., Weivoda, M.M., Inman, C.L., Ogrodnik, M.B., Hachfeld, C.M., Fraser, D.G., Onken, J.L., Johnson, K.O., Verzosa, G.C., Langhi, L.G.P., Weigl, M., Giorgadze, N., LeBrasseur, N.K., Miller, J.D., Jurk, D., Singh, R.J., Allison, D.B., Ejima, K., Hubbard, G.B., Ikeno, Y., Cubro, H., Garovic, V.D., Hou, X., Weroha, S.J., Robbins, P.D., Niedernhofer, L.J., Khosla, S., Tchkonia, T., Kirkland, J.L.: Senolytics improve physical function and increase lifespan in old age. Nat Med 24(8), 1246–1256 (2018). https://doi.org/10.1038/s41591-018-0092-9

[52] Koch, S., Larbi, A., Ozcelik, D., Solana, R., Gouttefangeas, C., Attig, S., Wikby, A., Strindhall, J., Franceschi, C., Pawelec, G.: Cytomegalovirus infection: a driving force in human t cell immunosenescence. Ann N Y Acad Sci 1114, 23–35 (2007). https://doi.org/10.1196/annals.1396.043

[53] Horvath, S., Levine, A.J.: Hiv-1 infection accelerates age according to the epigenetic clock. J Infect Dis 212(10), 1563–1573 (2015). https://doi.org/10.1093/infdis/jiv277

[54] Oltmanns, C., Liu, Z., Mischke, J., Tauwaldt, J., Mekonnen, Y.A., Urbanek-Quaing, M., Debarry, J., Maasoumy, B., Wedemeyer, H., Kraft, A.R.M., Xu, C.-J., Cornberg, M.: Reverse inflammaging: Long-term effects of hcv cure on biological age. J Hepatol 78(1), 90–98 (2023). https://doi.org/10.1016/j.jhep.2022.08.042

[55] Duncan, B.K., Miller, J.H.: Mutagenic deamination of cytosine residues in dna. Nature 287(5782), 560–561 (1980). https://doi.org/10.1038/ 287560a0

[56] Coulondre, C., Miller, J.H., Farabaugh, P.J., Gilbert, W.: Molecular basis of base substitution hotspots in escherichia coli. Nature 274(5673), 775– 780 (1978). https://doi.org/10.1038/274775a0

[57] Houseman, E.A., Accomando, W.P., Koestler, D.C., Christensen, B.C., Marsit, C.J., Nelson, H.H., Wiencke, J.K., Kelsey, K.T.: Dna methylation arrays as surrogate measures of cell mixture distribution. BMC Bioinformatics 13, 86 (2012). https://doi.org/10.1186/1471-2105-13-86

[58] Aryee, M.J., Jaffe, A.E., Corrada-Bravo, H., Ladd-Acosta, C., Feinberg, A.P., Hansen, K.D., Irizarry, R.A.: Minfi: a flexible and comprehensive bioconductor package for the analysis of infinium dna methylation microarrays. Bioinformatics 30(10), 1363–1369 (2014). https://doi.org/10.1093/bioinformatics/btu049

[59] Lehne, B., Drong, A.W., Loh, M., Zhang, W., Scott, W.R., Tan, S.-T., Afzal, U., Scott, J., Jarvelin, M.-R., Elliott, P., McCarthy, M.I., Kooner, J.S., Chambers, J.C.: A coherent approach for analysis of the illumina humanmethylation450 beadchip improves data quality and performance in epigenome-wide association studies. Genome Biol 16(1), 37 (2015). https://doi.org/10.1186/s13059-015-0600-x

[60] Benton, M.C., Johnstone, A., Eccles, D., Harmon, B., Hayes, M.T., Lea, R.A., Griffiths, L., Hoffman, E.P., Stubbs, R.S., Macartney-Coxson, D.: An analysis of dna methylation in human adipose tissue reveals differential modification of obesity genes before and after gastric bypass and weight loss. Genome Biol 16(1), 8 (2015). https://doi.org/10.1186/s13059-014-0569-x

[61] Fuchsberger, C., Abecasis, G.R., Hinds, D.A.: minimac2: faster genotype imputation. Bioinformatics 31(5), 782–784 (2015). https://doi.org/10.1093/bioinformatics/btu704

[62] Loh, P.-R., Danecek, P., Palamara, P.F., Fuchsberger, C., A Reshef, Y., K Finucane, H., Schoenherr, S., Forer, L., McCarthy, S., Abecasis, G.R., Durbin, R. L, Price, A.: Reference-based phasing using the haplotype reference consortium panel. Nat Genet 48(11), 1443–1448 (2016). https://doi.org/10.1038/ng.3679

[63] Das, S., Forer, L., Schönherr, S., Sidore, C., Locke, A.E., Kwong, A., Vrieze, S.I., Chew, E.Y., Levy, S., McGue, M., Schlessinger, D., Stambolian, D., Loh, P.-R., Iacono, W.G., Swaroop, A., Scott, L.J., Cucca, F., Kronenberg, F., Boehnke, M., Abecasis, G.R., Fuchsberger, C.: Next-generation genotype imputation service and methods. Nat Genet 48(10), 1284–1287 (2016). https://doi.org/10.1038/ng.3656

[64] McCarthy, S., Das, S., Kretzschmar, W., Delaneau, O., Wood, A.R., Teumer, A., Kang, H.M., Fuchsberger, C., Danecek, P., Sharp, K., Luo, Y., Sidore, C., Kwong, A., Timpson, N., Koskinen, S., Vrieze, S., Scott, L.J., Zhang, H., Mahajan, A., Veldink, J., Peters, U., Pato, C., van Duijn, C.M., Gillies, C.E., Gandin, I., Mezzavilla, M., Gilly, A., Cocca, M., Traglia, M., Angius, A., Barrett, J.C., Boomsma, D., Branham, K., Breen, G., Brummett, C.M., Busonero, F., Campbell, H., Chan, A., Chen, S., Chew, E., Collins, F.S., Corbin, L.J., Smith, G.D., Dedoussis, G., Dorr, M., Farmaki, A.-E., Ferrucci, L., Forer, L., Fraser, R.M., Gabriel, S., Levy, S., Groop, L., Harrison, T., Hattersley, A., Holmen, O.L., Hveem, K., Kretzler, M., Lee, J.C., McGue, M., Meitinger, T., Melzer, D., Min, J.L., Mohlke, K.L., Vincent, J.B., Nauck, M., Nickerson, D., Palotie, A., Pato, M., Pirastu, N., McInnis, M., Richards, J.B., Sala, C., Salomaa, V., Schlessinger, D., Schoenherr, S., Slagboom, P.E., Small, K., Spector, T., Stambolian, D., Tuke, M., Tuomilehto, J., Van den Berg, L.H., Van Rheenen, W., Volker, U., Wijmenga, C., Toniolo, D., Zeggini, E., Gasparini, P., Sampson, M.G., Wilson, J.F., Frayling, T., de Bakker, P.I.W., Swertz, M.A., McCarroll, S., Kooperberg, C., Dekker, A., Altshuler, D., Willer, C., Iacono, W., Ripatti, S., Soranzo, N., Walter, K., Swaroop, A., Cucca, F., Anderson, C.A., Myers, R.M., Boehnke, M., McCarthy, M.I., Durbin, R.: A reference panel of 64,976 haplotypes for genotype imputation. Nat Genet 48(10), 1279–1283 (2016). https://doi.org/10.1038/ng.3643

[65] Delaneau, O., Ongen, H., Brown, A.A., Fort, A., Panousis, N.I., Dermitzakis, E.T.: A complete tool set for molecular qtl discovery and analysis. Nat Commun 8, 15452 (2017). https://doi.org/10.1038/ncomms15452

[66] Ernst, J., Kellis, M.: Chromhmm: automating chromatin-state discovery and characterization. Nat Methods 9(3), 215–216 (2012). https://doi.org/10.1038/nmeth.1906

[67] Elsworth, B., Lyon, M., Alexander, T., Liu, Y., Matthews, P., Hallett, J., Bates, P., Palmer, T., Haberland, V., Smith, G.D., Zheng, J., Haycock, P., Gaunt, T.R., Hemani, G.: The mrc ieu opengwas data infrastructure. bioRxiv (2020) https://www.biorxiv.org/content/early/2020/08/10/2020.08.10.244293.full.pdf. https://doi.org/10.1101/2020.08.10.244293

